# Sex differences in the genetic basis of human recombination within 190,000 parent-child pairs

**DOI:** 10.64898/2026.04.30.721836

**Authors:** Ilse Krätschmer, Laura Hegemann, Gunnar Pálsson, Robin Hofmeister, Anne Richmond, Caroline Hayward, Ole A. Andreassen, Astanand Jugessur, Ida Elken Sønderby, Adrian Dahl Askelund, Estonian Biobank Research Team, Zoltan Kutalik, Riccardo E. Marioni, Marteinn Thor Hardarson, Ólafur A. Stefansson, Hákon Jónsson, Peter Kharchenko, Kári Stefánsson, Bjarni V. Halldórsson, Alexandra Havdahl, Matthew R. Robinson

## Abstract

**Introduction:** Sexual reproduction uses a specialized cell division called meiosis, in which a single round of DNA replication is followed by two cell divisions to create haploid gametes. Genetic recombination in meiosis assures faithful segregation of chromosomes and establishes patterns of genetic linkage and inheritance. Meiotic recombination is thus a fundamental genomic process that shapes major features of the genomic landscape, influences mutation, and creates genetic diversity.

**Rationale:** The genetic basis of between-individual variation in the occurrence and genomic location of meiotic recombination events remains poorly understood in humans. In particular, we lack an understanding of whether the same DNA regions shape variation in recombination rates across studies. We address this by curating 112,144 maternal and 78,653 paternal meiosis events (parent-child pairs) across cohorts from four countries, creating the largest dataset of its kind.

**Results:** We identify 58 independent genomic regions associated with the occurrence and genomic location of recombination events, 24 of which are unique across phenotypes. Of the 24 unique regions, 6 are novel and specific to female meiotic recombination harboring genes linked to ovarian function, a 7th novel region is linked to male recombination. Many regions have significant sex-dependent effects across studies. We estimate the between-sex genetic correlation for meiotic recombination rate to be 0.374 (0.072 SD), which captures the extent to which the genetic factors influencing a trait in one sex also impact the same trait in the other. 30-40% of the individual-level variation in recombination rate is attributable to DNA markers, of which only 37.7% (2.7% SD) and 47.1% (3.8% SD) is attributable to previously known global modulators of meiotic recombination, in females and males respectively. We create a polygenic predictor that significantly predicts recombination rates of both women and men after adjusting for age and year of birth. We find significant correlations between female meiotic recombination, reproductive outcomes and reproductive aging, after adjusting for study ascertainment.

**Conclusions:** In the largest meta-analysis to date of human meiotic recombination, we identify novel loci with sex-specific effects that influence male and female recombination rate across multiple cohorts.

---

In humans and other organisms, meiosis differs across the biological sexes, both in terms of timing and in the number of gametes produced (*1–5*). Meiotic recombination, the exchange of DNA between homologous chromosomes through chromosomal crossover and gene-conversion events, enhances genetic diversity and plays a fundamental role in ensuring the proper segregation of chromosomes (*6*). The biological sexes differ in the rate of meiotic recombination and in the distribution of recombination events throughout the genome. While female autosomal recombination rates are higher than those of males over the majority of the genome in humans, male recombination is elevated towards the telomeres (*1*). Within each sex, there is also between-individual variation in the occurrence and genomic location of recombination events (*5, 7–10*) and recent studies suggest that DNA regions shape variation in recombination rates across the biological sexes in different ways (*10*). However, studies of the genetic basis of meiotic recombination are generally in single cohorts of limited sample size and lack replication.

Here, we combine genetic data of parents and children from four Northern European biobanks. For three we have individual-level data: Norway (Norwegian Mother, Father and Child Cohort Study, MoBa (*11*)), Estonia (Estonian Biobank, EstBB), and Scotland (Generation Scotland, GS). Sample sizes for each study are given in Table S1. We infer autosomal double strand crossover positions in the gametes transmitted from each parent to their offspring using a novel state-of-the-art Identity-By-Descent (IBD) mapping approach, modified from Ref. (*12*). For each mother-child pair, we obtain an observation of female meiotic recombination and an observation of paternal meiotic recombination for each father-child pair. For each female and male meiosis, we record four different values: (i) the total number of double-strand break cross-overs occurring on the autosomes (termed CO throughout); (ii) the proportion of the gamete’s paternally or maternally derived genome that is from a single grand-parental origin (PA), which is a measure of the probability that the alleles of a randomly chosen pair of loci are shuffled during formation of the gamete (equivalently, the proportion of locus pairs at which the alleles are shuffled, referred to in the literature as 𝑟_𝑖𝑛𝑡𝑟𝑎_ (*13*)); (iii) the mean distance (in bp) between consecutive autosomal cross-over events (MD), which is a proxy for interference; and (iv) the average relative distance of the closest crossovers from the telomere across chromosomes in cM (telomere distance, TD).

We find that the four phenotypes differ among the biological sexes in a consistent manner across the three cohorts (Figure 1, Table S1). Figure 1 shows the similarity of the normalized distribution of the four recombination phenotypes within each sex across countries, despite each of the cohorts having different relatedness structures and different single-nucleotide polymorphism (SNP) array genotyping technologies. Additionally, the phenotypic relationships among the meiotic recombination phenotypes within females and males are also the same across countries, as shown in Figure S1A.

**Figure 1:**
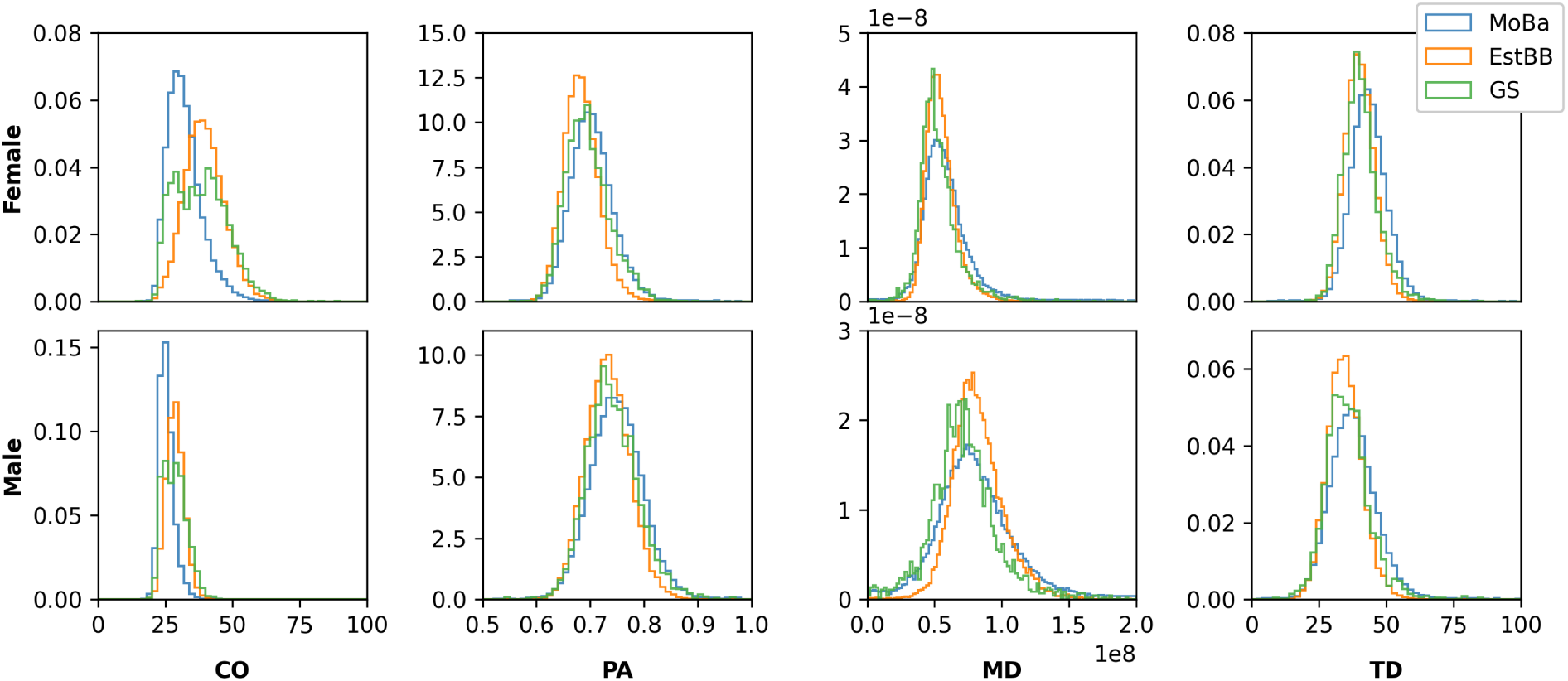
Density plots of the distribution of four recombination phenotypes. Distribution of female (top) and male (bottom) meioses for recombination cross-over rate (CO), the proportion of single transferred ancestry (PA), the mean distance among consecutive cross-over events (MD), and the telomere distance (TD) across three different European study cohorts (MoBa, EstBB and GS).

We then combine these data with previously published data from Iceland (*10*) (deCODE Genetics), where the four phenotypes were recalculated for 41,745 women and 30,184 men. This gives a total sample size of 112,144 maternal meioses events (mother-child pairs) for 82,500 mothers and 78,653 paternal meioses (father-child pairs) for 62,500 fathers, which is far larger than the largest previous genome scan of variants influencing genome-wide recombination rate that was conducted in a single cohort only, where replication was not attempted within another study (*10*).

## Novel genetic loci for meiotic recombination with sex-specific effects

For each biological sex, we estimate ordinary least squares (OLS) regression coefficients common to genome-wide association studies (GWAS), which quantify the marginal relationship of ∼ 9M imputed common SNP variants to each of the four recombination phenotypes. The OLS GWAS estimates for each SNP are calculated separately within EstBB, MoBa and Iceland cohorts, using a leave-one-chromosome-out (LOCO) mixed linear model approach. These are then meta-analyzed together within a fixed-effect meta analysis framework (see Methods). We conservatively define associated regions by taking a focal lead variant (that of strongest association) and fitting a joint model within a 10Mb window around the variant (using the default parameters of GCTA’s cojo algorithm). Thus, significant associated variants ≥ 10Mb from each other are treated as independent. We find 58 genomic regions associated with the four recombination characteristics at a genome-wide significance threshold of 5·10^−8^, of which 24 are unique across phenotypes (Table 1). Table S2 gives the number of SNPs passing the genome-wide significance threshold within each genomic region found here. Table S3 gives the number of SNPs fit jointly for both sexes and passing a genome-wide significance threshold of 5 · 10^−8^ within each region. We also perform fine-mapping with SuSiEx (*14*) across men and women within each region, to identify credible sets of SNPs that are shared or specific to each sex, as those with a posterior inclusion probability ≥ 0.9 of being associated in either men, women, or both (Table S4). A full list of genes that are mapped to leading SNPs in significantly associated genomic regions using FUMA (*15*) are given in Tables S5-S8 for the four phenotypes. A visualization of the statistical significance of the meta-analyzed effects for the four phenotypes in men and women is given in Figure 2. Quantile-quantile plots, shown in Figure S2, and the SumHer intercept values, given in Table S9, suggest that relatedness, repeated measures and potential confounding within the data are well controlled. We also find that our significant loci replicate strongly within the held-out Scottish GS cohort (Figure S3).

**Figure 2:**
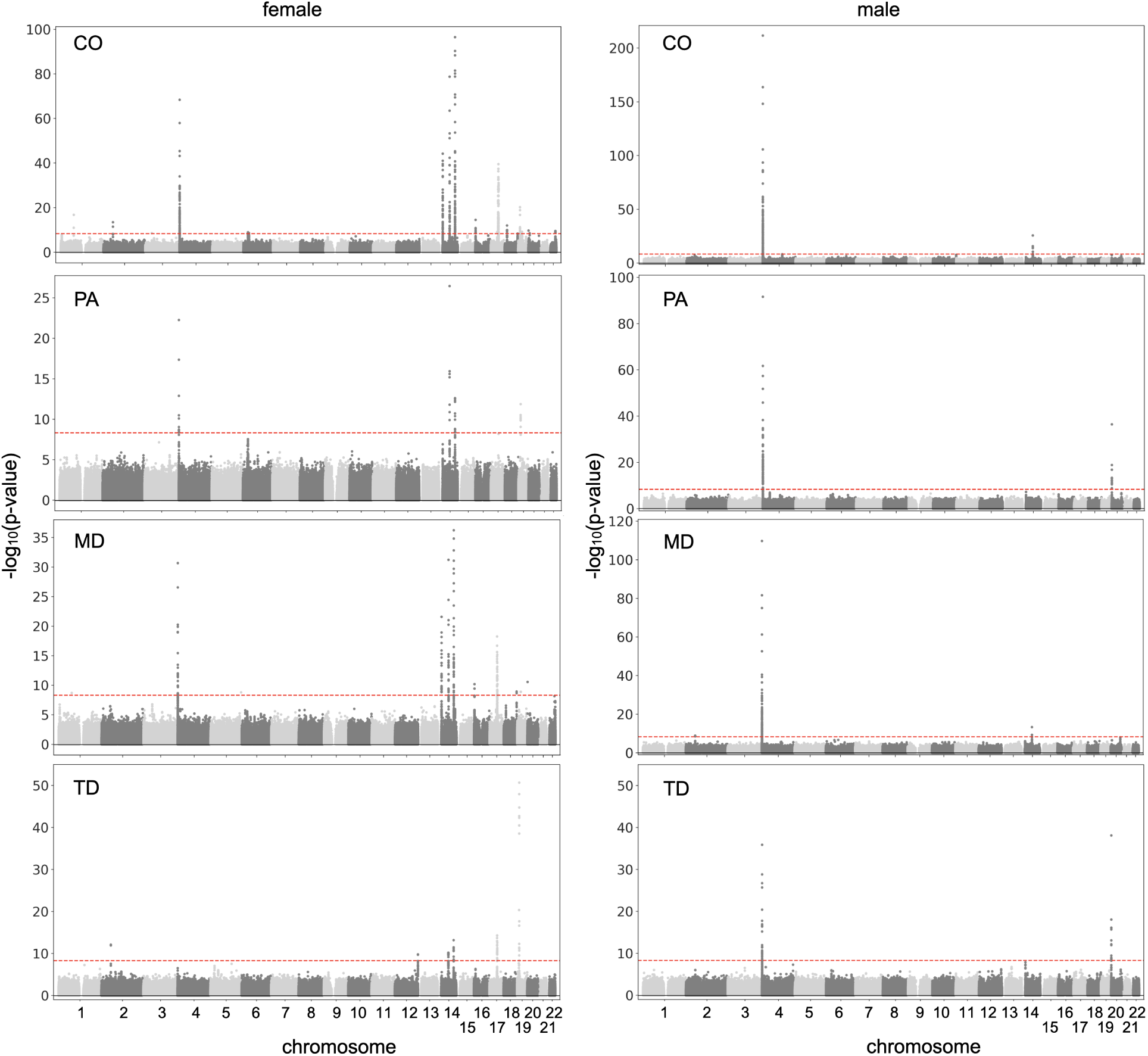
Manhattan plots for four recombination phenotypes of female and male meioses. Phenotypes analyzed are recombination cross-over rate (CO), the proportion of single transferred ancestry (PA), the mean distance among consecutive cross-over events (MD), and the telomere distance (TD). Each point depicts the −𝑙𝑜𝑔_10_ of the p-value for each SNP obtained from a meta-analysis across cohorts. The dashed red line represents the multiple testing-corrected p-value threshold of 5 · 10^−8^. Estimates for female recombination phenotypes are given in the left column and those for males are given in the right column.

**Table 1:**
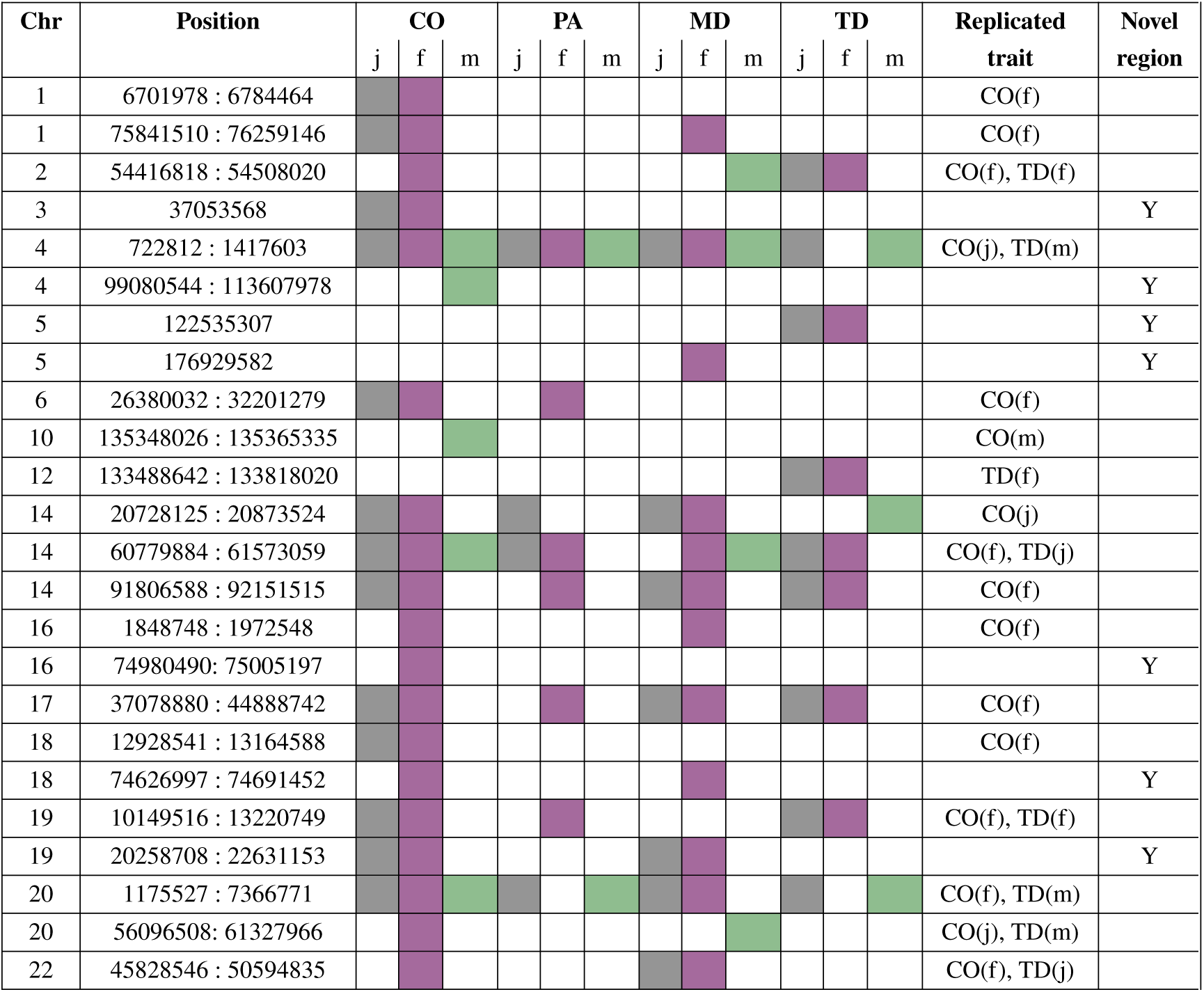
Independent DNA regions containing genome-wide significantly associated loci. The association analyses are performed separately in females (f) and males (m) and jointly (j) for both sexes. Four traits are considered: recombination rate (CO), the proportion of single transferred ancestry (PA), the telomere distance (TD) and the mean distance among consecutive cross-over events (MD). For each trait and analysis, colored squares indicate that at least one genome-wide significant locus is discovered for a certain region given by chromosome (chr) and base pair window (position). The column ’Replicated trait’ indicates replication of the region found in Ref. (*10*). ’Y’ in the column ’Novel region’ indicates that a region was not previously reported as associated with any of the four phenotypic phenotypes.

We find seven novel regions which are summarized in Table 2. Table 1 provides an indicator of whether each of the 24 DNA regions contains SNPs that were previously discovered in the most recent GWAS of human recombination that was conducted within Iceland (*10*). Ref. (*10*) report 35 jointly fit SNPs, within 18 independent genomic regions (≥ 2Mb from any other) without an attempt of replication within another study. Here, we find that 29 of the 35 previously reported SNPs replicate at genome-wide significance within our meta-analyses, within 17 independent 10Mb genomic regions (Table 1). Of the six SNPs found within the most recent Iceland study (*10*) which do not replicate here (and are ≥ 10Mb away from any significant association), one is associated with a phenotype not considered here (see Supplementary Text), and another three are not available in MoBa and EstBB after quality control. Of the remaining two, rs6889665 has the same sign in both MoBa and EstBB with a p-value of 3.23 · 10^−7^ in EstBB, which confirms a possible association, but does not pass the genome-wide significance threshold used in our meta-analysis. The last SNP rs11165778 has the same sign in MoBa and EstBB, but is below the significance threshold (see Supplementary Table S10). Our results also replicate all regions identified in other smaller scale GWAS analyses of meiotic recombination that have been conducted to date (*5, 8, 16*) and a recent retrospective analysis of pre-implantation genetic testing data whose variants are significantly associated with recombination rate and meiotic aneuploidy risk (*17*).

**Table 2:**
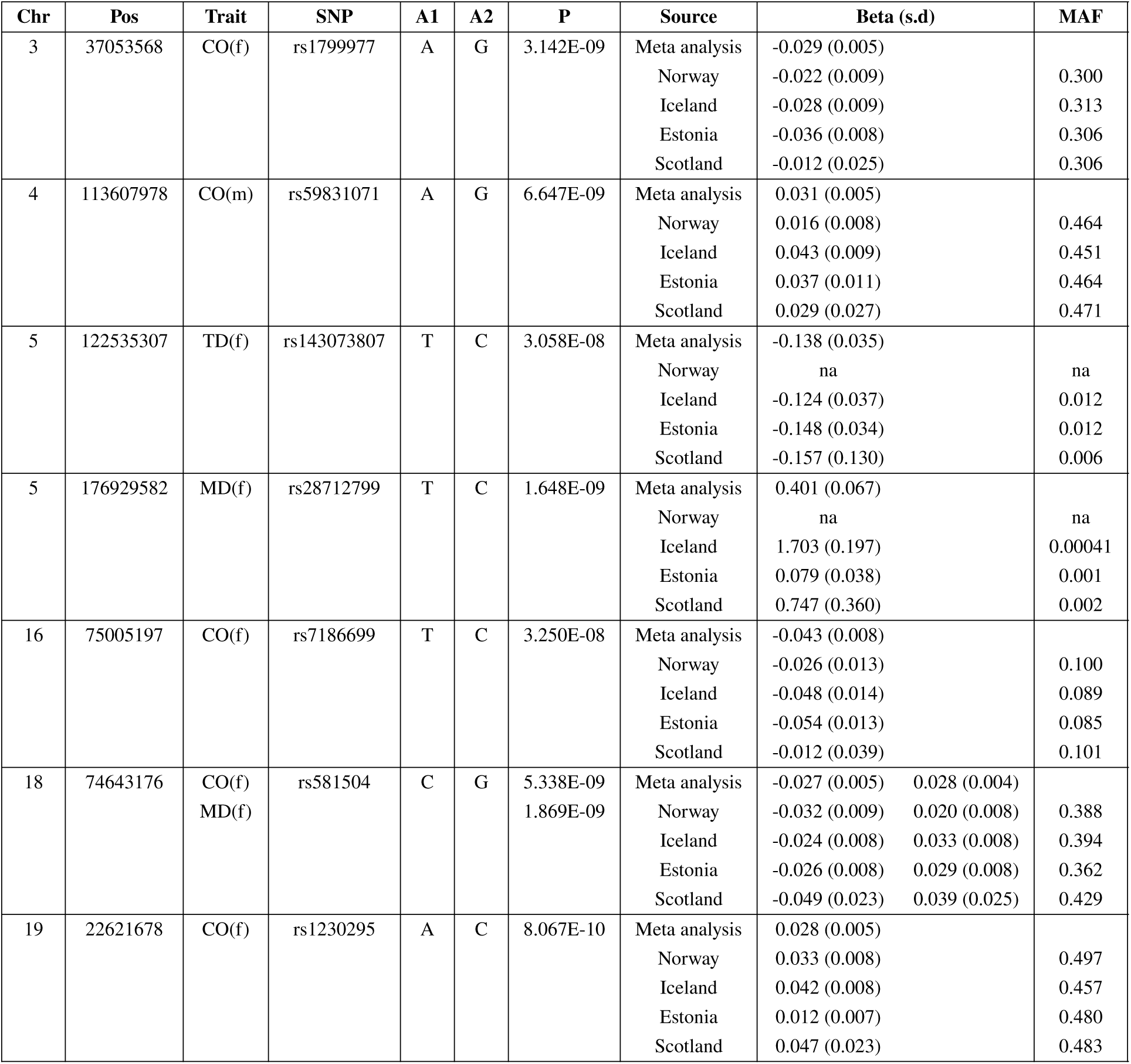
Novel independent DNA regions containing genome-wide significantly associated loci. The association analyses are performed separately in females (f) and males (m) for four traits: recombination rate (CO), the proportion of single transferred ancestry (PA), the telomere distance (TD) and the mean distance among consecutive cross-over events (MD). For each region identified by chromosome (Chr) and base pair window (Pos), the leading single nucleotide polymorphism (SNP), the two alleles (A1, A2), p-value (P) form the meta-analysis, the effect sizes (Beta) and minor allele frequency (MAF) are given for the different cohorts and meta-analysis (Source). ’na’ indicates that the SNP was not present after QC in the cohort individuals.

Table 2 gives the focal lead variant, the genomic locations, significance, and the effect sizes and frequencies across countries of the seven novel regions we discover, which contain genes that to our knowledge have not previously been explicitly implicated in human meiotic recombination, and which replicate across at least two cohorts. Of the seven novel regions we discover, six are specific to female meiotic recombination phenotypes. First, chromosome 3 bp region 37053568 containing rs1799977 is associated with female CO, which is a missense variant in *MLH1*, a DNA mismatch repair protein. *MLH1* has been shown to provide instructions for DNA repair proteins which fix errors during DNA replication in mice and yeast (*18*). Mutations in *MLH1* are also linked to Lynch syndrome, a hereditary cancer predisposition, through DNA mismatch repair system (MMR) that fixes replication errors outside of meiosis (*19*).

Second, chromosome 5 bp 122535307 contains rs143073807, an intergenic SNP associated with telomere distance in women that is 400kb from *PRDM6*. *PRDM*s are able to recruit histone ‘writers’ and ‘erasers’ (i.e., demethylases and deacetylases), including *PRDM6*, which is a poorly characterized member of the family (*20*). It has been proposed that *PRDM6* is a transcriptional repressor, particularly within the cardiovascular system (*20*). There is no evidence of direct DNA binding by *PRDM6* or a role in meiosis, but it is predicted to recognize a G/T-rich motif with its four *ZNF*s (*20*). *PRDM6* is a gene that is expressed in oocytes in pigs and potentially humans (expressed in ovaries in GTEX data), where it is believed to function as a maternal transcript involved in early embryo development and epigenetic reprogramming (*21*). However, this association is fine-mapped by SuSiEx to a higher base pair position of 123614260 at SNP rs31319, which is an intronic variant in long intergenic non-protein coding RNA 1170 (*LINC01170*). *LINC01170* may be linked to complex chromosomal rearrangements and promote open chromatin so that the REC114-MEI4-IHO1 complex can more easily bind to the DNA, ensuring the REC114 complex attaches to the correct ”hotspots” rather than random sites across the genome.

Third, chromosome 5 bp 176929582, is associated with female MD, with SNP rs28712799 as the most strongly associated, which is a 3’ UTR variant in Docking Protein 3 (*DOK3*). *DOK3* has not been identified as a primary meiotic recombination gene, but it has been linked to genomic instability and DNA repair processes that are critical for proper recombination, with *DOK3* expression correlating with *MLH1* and *MSH2* that are key genes involved in meiotic pathways and DNA damage response (*22, 23*). The next closest gene highlighted by FUMA is *DDX41*, which is crucial for viability in mice and humans, with complete loss being embryonically lethal (*24*). The specific direct role of *DDX41* in human meiosis has not been characterized, but in the roundworm *Caenorhabditis elegans*, the *DDX41* homolog, *sacy-1*, is identified as a factor essential for oocyte meiotic maturation, where an oocyte resumes and completes meiosis (*25*). The proposed biological functions of nuclear *DDX41* include R-loop resolution, small nucleolar RNA processing and ribosomal RNA (rRNA) processing. *DDX41* is associated with hematopoietic stem cell proliferation and differentiation (*26*). SuSiEx fine-maps the association to a lower base pair position of 173976464, an intronic and upstream variant of *LOC105377739*, an uncharacterized non-coding RNA. Thus, there many potential candidate genes within this region.

Fourth, chromosome 16 bp region 74980490–75005197 containing rs7186699 is an intronic variant of *WDR59*. In Drospohila and yeast, *WDR59* functions in the GATOR-TORC1 signaling pathway and has been linked to the regulation of events during oogenesis that are essential for successful meiotic progression, as *WDR59* depletion causes low *TORC1* activity, disrupting meiotic cycle maintenance (*27*). SuSiEx fine-maps the association to intronic variants rs143068306 and rs143235261 in Pyruvate Dehydrogenase Phosphatase Regulatory Subunit 2, (pseudogene) (*PDPR2P*). Both *WDR59* and *PDPR2P* shared the same promoters and enhancers are located in the 16p13.3 region, which is prone to copy number variations and rearrangements and are proximal to genes that are known to be critical for meiosis, such as *REC114*.

Fifth, chromosome 18 bp region 74626997–74691452 is associated with female CO and MD, at GWAS lead SNP rs581504, an intronic variant of Zinc Finger Protein 236 (*ZNF236*) and is fine-mapped by SuSiEx to intronic variants rs4640270 and rs8084848 of neighboring *ZNF236-DT*. *ZNF236* is within a region prone to copy number variation and is associated with Chromosome 18Q Deletion Syndrome. While not primarily known as a core meiotic recombination protein, it is listed alongside meiotic and DNA repair genes in specific high-throughput screenings in *Drospohila*, where a homologous *C2H2* zinc finger protein is associated with the induction of double strand breaks in female meiosis (*28*).

Sixth, chromosome 19 bp region 20258708–22631153, associated with female CO and MD, is a region of the genome known for having an unusually high average recombination rate and zinc-finger (*ZNF*) gene clusters. It has lead GWAS (rs230295) and fine-mapped (rs12977311) variants upstream and is intronic to Zinc Finger Protein 98 (*ZNF98*). *ZNF98* is a C2H2 zinc finger protein, a family often involved in transcriptional regulation, but also implicated in chromatin modification and DNA interaction during meiosis (*29*). The top STRING interactants also contain Cohesin subunit SA-2, a component of the cohesin complex, which is required for the cohesion of sister chromatids after DNA replication. Although there is no established direct role in meiotic recombination, our findings of two associated C2H2 zinc finger proteins suggest a prominent role of this largest but poorly characterized family of transcription factors, which is supported by studies in non-human species (*30, 31*).

Finally, we find a single novel region for male recombination: chromosome 4 bp region 99080544–113607978, which SuSiEx fine-maps to rs973126, a synonymous variant in bifunctional 3’-phosphoadenosine 5’-phosphosulfate synthase 1 (*PAPSS1*). *PAPSS1* plays a crucial role in mitigating DNA damage or promoting efficient DNA repair. Its suppression leads to increased DNA double-strand breaks (*32*), which is thought to result from impaired DNA repair mechanisms or increased accumulation of DNA damage, possibly mediated by impaired post-translational sulfation of nuclear proteins crucial for repairing DNA lesions. *PAPSS1* expression is documented in human testis tissue, specifically during spermatogenesis. Sulfation is crucial for the biosynthesis of seminolipid, a sulfoglycolipid essential for male fertility.

For the regions we identify as significantly associated with female and male meiotic recombination, we test for sex differences in the effect sizes of the associated loci by: *(i)* testing for a z-score difference in the meta-analysis effect sizes between the biological sexes for each SNP genome-wide; and *(ii)* by taking the focal variants of greatest association within each region and testing for a SNP-by-sex interaction using a mixed-effects model within the three cohorts, MoBa, EstBB and GS. The Scottish cohort, GS, was not included in the meta-analysis (Table 3 and see Methods). Of the 21 regions identified for CO, eight show loci with a genome-wide significant difference in SNP effect size between the biological sexes, using the z-score difference test. Six out of the seven regions identified for PA, six of 14 regions identified for MD, and five out of ten regions identified for TD show multiple loci with a genome-wide significant difference in the SNP effect size between the biological sexes. As shown in Tables S3 and S4, many regions contain multiple variants, multiple credible sets and multiple SNPs within credible sets. This may reflect either multiple underlying causal variants, or multiple associated variants that capture a single underlying association. We cannot distinguish among these or resolve them without full whole genome sequence data for all cohorts and larger sample sizes. But we provide evidence for a range of regions, from those whose marginal effect sizes are consistent, to those which differ significantly among the biological sexes.

**Table 3:**
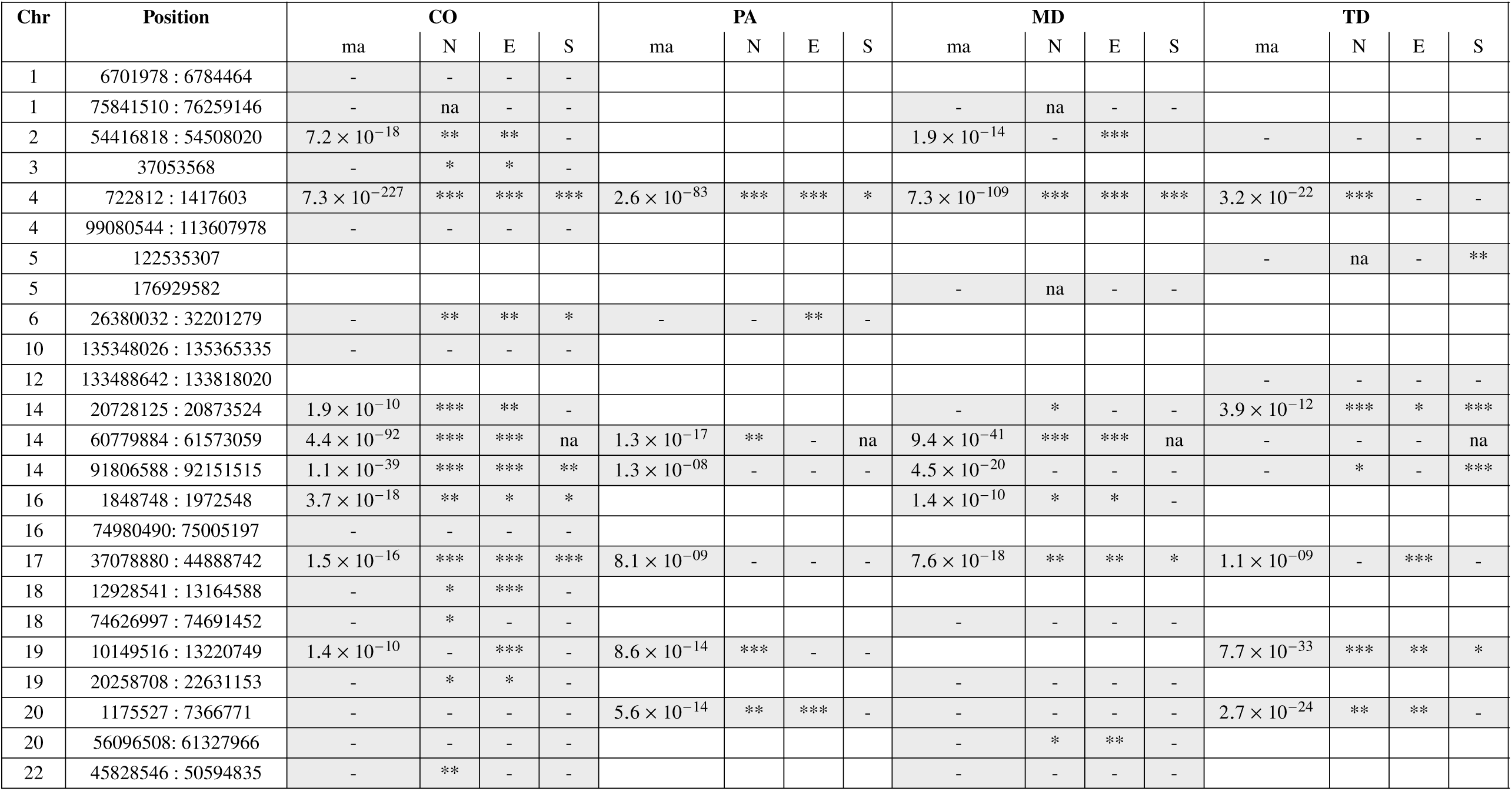
Interaction of loci and biological sex for genome-wide significantly associated loci. Four traits are considered: recombination rate (CO), the proportion of single transferred ancestry (PA), the telomere distance (TD) and the mean distance among consecutive cross-over events (MD). For each genomic region indicated by chromosome (Chr) and base pair window (Pos), the column ’ma’ gives the minimum p-value for SNPs within each region from a test of the difference in effect sizes between female and male recombination phenotypes with ’-’ indicating a p-value above 5 · 10^−8^. Within-study interaction testing was also conducted for Norway (N), Estonia (E), and Scotland (S). Significance of within-study interaction testing: ’-’ refers to a p-value above 0.05; ’*’ p-value between 0.01 and 0.05; ’**’ to p-value between 0.01 and 0.001; ’***’ to p-value below 0.001; ’na’ indicates that SNPs in the region do not pass imputation QC. Grey boxes indicate traits and countries for which within-study interaction testing is conducted.

| Chr | Position | CO |  |  |  | PA |  |  |  | MD |  |  |  | TD |  |  |  |
| --- | --- | --- | --- | --- | --- | --- | --- | --- | --- | --- | --- | --- | --- | --- | --- | --- | --- |
|  |  | ma | N | E | S | ma | N | E | S | ma | N | E | S | ma | N | E | S |
| 1 | 6701978 : 6784464 | - | - | - | - |  |  |  |  |  |  |  |  |  |  |  |  |
| 1 | 75841510 : 76259146 | - | na | - | - |  |  |  |  | - | na | - | - |  |  |  |  |
| 2 | 54416818 : 54508020 | $7.2 \times 10^{-18}$ | ** | ** | - | | | | | $1.9 \times 10^{-14}$ | - | *** | | - | - | - | - |
| 3 | 37053568 | - | * | * | - |  |  |  |  |  |  |  |  |  |  |  |  |
| 4 | 722812 : 1417603 | $7.3 \times 10^{-227}$ | *** | *** | *** | $2.6 \times 10^{-83}$ | *** | *** | * | $7.3 \times 10^{-109}$ | *** | *** | *** | $3.2 \times 10^{-22}$ | *** | - | - |
| 4 | 99080544 : 113607978 | - | - | - | - |  |  |  |  |  |  |  |  |  |  |  |  |
| 5 | 122535307 |  |  |  |  |  |  |  |  |  |  |  |  | - | na | - | ** |
| 5 | 176929582 |  |  |  |  |  |  |  |  | - | na | - | - |  |  |  |  |
| 6 | 26380032 : 32201279 | - | ** | ** | * | - | - | ** | - |  |  |  |  |  |  |  |  |
| 10 | 135348026 : 135365335 | - | - | - | - |  |  |  |  |  |  |  |  |  |  |  |  |
| 12 | 133488642 : 133818020 |  |  |  |  |  |  |  |  |  |  |  |  | - | - | - | - |
| 14 | 20728125 : 20873524 | $1.9 \times 10^{-10}$ | *** | ** | - | | | | | - | * | - | - | $3.9 \times 10^{-12}$ | *** | * | *** |
| 14 | 60779884 : 61573059 | $4.4 \times 10^{-92}$ | *** | *** | na | $1.3 \times 10^{-17}$ | ** | - | na | $9.4 \times 10^{-41}$ | *** | *** | na | - | - | - | na |
| 14 | 91806588 : 92151515 | $1.1 \times 10^{-39}$ | *** | *** | ** | $1.3 \times 10^{-08}$ | - | - | - | $4.5 \times 10^{-20}$ | - | - | - | - | * | - | *** |
| 16 | 1848748 : 1972548 | $3.7 \times 10^{-18}$ | ** | * | * | | | | | $1.4 \times 10^{-10}$ | * | * | - | | | | |
| 16 | 74980490 : 75005197 | - | - | - | - |  |  |  |  |  |  |  |  |  |  |  |  |
| 17 | 37078880 : 44888742 | $1.5 \times 10^{-16}$ | *** | *** | *** | $8.1 \times 10^{-09}$ | - | - | - | $7.6 \times 10^{-18}$ | ** | ** | * | $1.1 \times 10^{-09}$ | - | *** | - |
| 18 | 12928541 : 13164588 | - | * | *** | - |  |  |  |  |  |  |  |  |  |  |  |  |
| 18 | 74626997 : 74691452 | - | * | - | - |  |  |  |  | - | - | - | - |  |  |  |  |
| 19 | 10149516 : 13220749 | $1.4 \times 10^{-10}$ | - | *** | - | $8.6 \times 10^{-14}$ | *** | - | - | | | | | $7.7 \times 10^{-33}$ | *** | ** | * |
| 19 | 20258708 : 22631153 | - | * | * | - |  |  |  |  | - | - | - | - |  |  |  |  |
| 20 | 1175527 : 7366771 | - | - | - | - | $5.6 \times 10^{-14}$ | ** | *** | - | - | - | - | - | $2.7 \times 10^{-24}$ | ** | ** | - |
| 20 | 56096508 : 61327966 | - | - | - | - |  |  |  |  | - | * | ** | - |  |  |  |  |
| 22 | 45828546 : 50594835 | - | ** | - | - |  |  |  |  | - | - | - | - |  |  |  |  |

## Variation of meiotic recombination shared between the biological sexes

We estimate the proportion of variation within each sex attributable to common genetic variants genome-wide (termed the SNP-heritability, 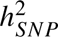) and the proportion of variance that is shared between the two sexes (between-sex genetic correlation). We do this in two ways: *(i)* by applying an adapted version of a multivariate Bayesian multiple linear regression model, MAJA (*33*), using individual-level data of more than 1M SNPs from EstBB and MoBa separately; and *(ii)* by using our GWAS effect size estimates from EstBB, MoBa, deCODE and the meta-analysis to calculate 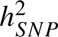 with a Bayesian summary statistic linear penalized regression model (labeled SBayesRC (*34*)) and a frequentist LD score-based approach (labeled SumHer (*35*)). More details on the statistical frameworks can be found in Methods.

We find consistent estimates of 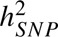 across the three cohorts, and similar estimates across the sexes and across different inference methods, as shown in Figure 3A. We find higher 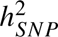 estimates for CO than for the other traits, with female estimates being higher than those for males. For the traits of lower 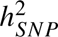, we observe that SBayesRC and SumHer frequently give estimates that are not different to zero within at least one of the cohorts reflecting a lack of power.

**Figure 3:**
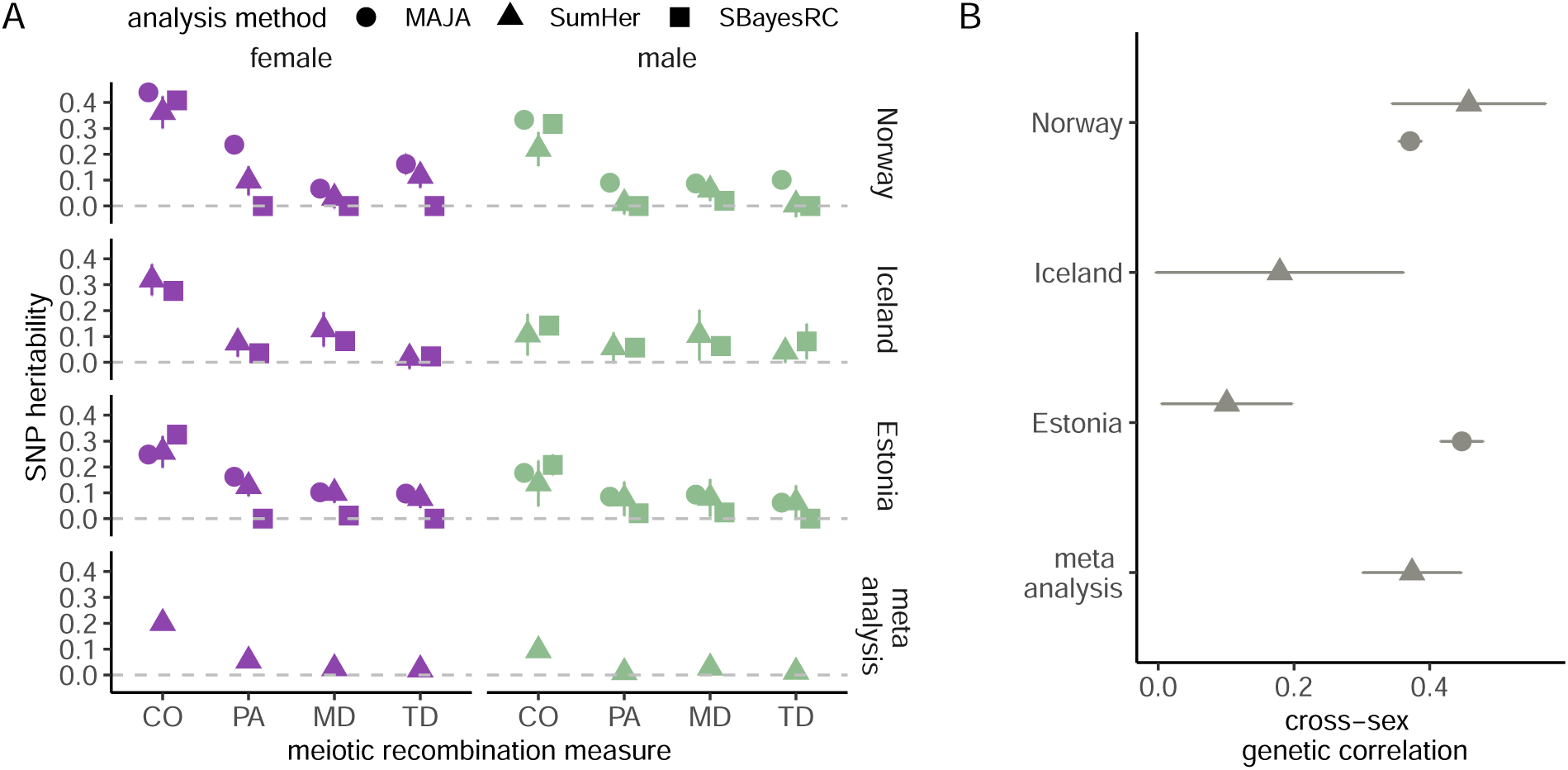
The proportion of phenotypic variance attributable to common genetic variation for four phenotypes of meiotic recombination and the cross-sex genetic correlation for recombination rate, across data from three European countries. In (**A**), the proportion of trait variation attributable to common single nucleotide polymorphisms (SNP heritability) estimated by three statistical approaches: a Bayesian multiple-outcome linear penalized regression model applied to individual-level data (labeled MAJA), a Bayesian linear penalized regression model applied to summary statistic association estimates (labeled SBayesRC), and a frequentist LD Score-based approach applied to summary statistic association estimates (labeled SumHer). Four recombination phenotypes of female and male meioses are analyzed: recombination cross-over rate (CO), the proportion of single transferred ancestry (PA), the mean distance among consecutive cross-over events (MD), and the telomere distance (TD). For CO, which shows the highest SNP heritability across countries for both female and male phenotypes, (**B**) shows the between-sex genetic correlation across studies. Note that SumHer estimates in Estonia are underpowered because of the small sample size for males compared to females. Error bars give the 95% intervals of the estimates. Where values are missing, individual-level data was not accessed (Iceland). For more details on the used statistical frameworks, see Methods.

For CO, where 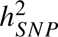 is significantly different from zero for both men and women across all cohorts and inference approaches, we proceed to estimate the between-sex genetic correlation, which captures the extent to which the genetic factors influencing a trait in one sex also affect the same trait in the other sex. We find that estimates range from 0.1 to 0.45, as shown in Figure 3B. In MoBa, where summary statistic estimates are well-powered and individual-level estimates are obtained, correlation estimates are consistent across inference methods. These are also consistent with individual-level estimates from EstBB and the estimate made from meta-analysis values. Phenotypic and genetic correlations among the four recombination phenotypes across studies and between sex-genetic correlations for all recombination phenotypes are given in Supplementary Figures S1 and S4.

Previous studies (e.g. (*7, 36–38*)) have identified key processes and genes which play a mechanistic role in meiotic recombination. Previous work (*10*) has also shown that double strand breaks are more likely on average to be located in genomic regions with distinct characteristics. We ask a novel question: whether the genetic variants that influence differences among people in the rate and placement of recombination events also have specific characteristics. Using SBayesRC and the GWAS meta-analysis results, we partition the 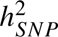 for CO across the genome for males and females across different genomic annotations.

We first calibrate data across published studies that have identified genomic regions with H3K4me3 and H3K27me3 epigenetic marks, as well as ATAC-seq ”tracks”, within both spermatocytes (pachytene stage) and oocytes. Crossover hotspots have been shown to be associated with euchromatin regions having specific chromatin modifications (*36*). Here, we map SNPs to these features to infer the degree to which population-level variation within each sex across cohorts can be attributed to local sex-cell specific epigenetic and open chromatin regions. The results are shown in Figure 4. We find that only SNPs within H3K27me3 chromatin modifications contribute to the 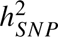 in males and females. SNPs within H3K27me3 chromatin modifications specific to oocytes make a proportional contribution of 3.8% to CO 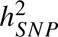 in females, with posterior probability ≥ 0.95 of the value being larger than expected given the number of SNPs mapped to the annotation. SNPs within H3K27me3 chromatin modifications specific to spermatocytes (pachytene stage) contribute 4.8% to CO 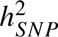 in males, with posterior probability ≥ 0.95 of the value being larger than expected given the number of SNPs mapped to the annotation.

**Figure 4:**
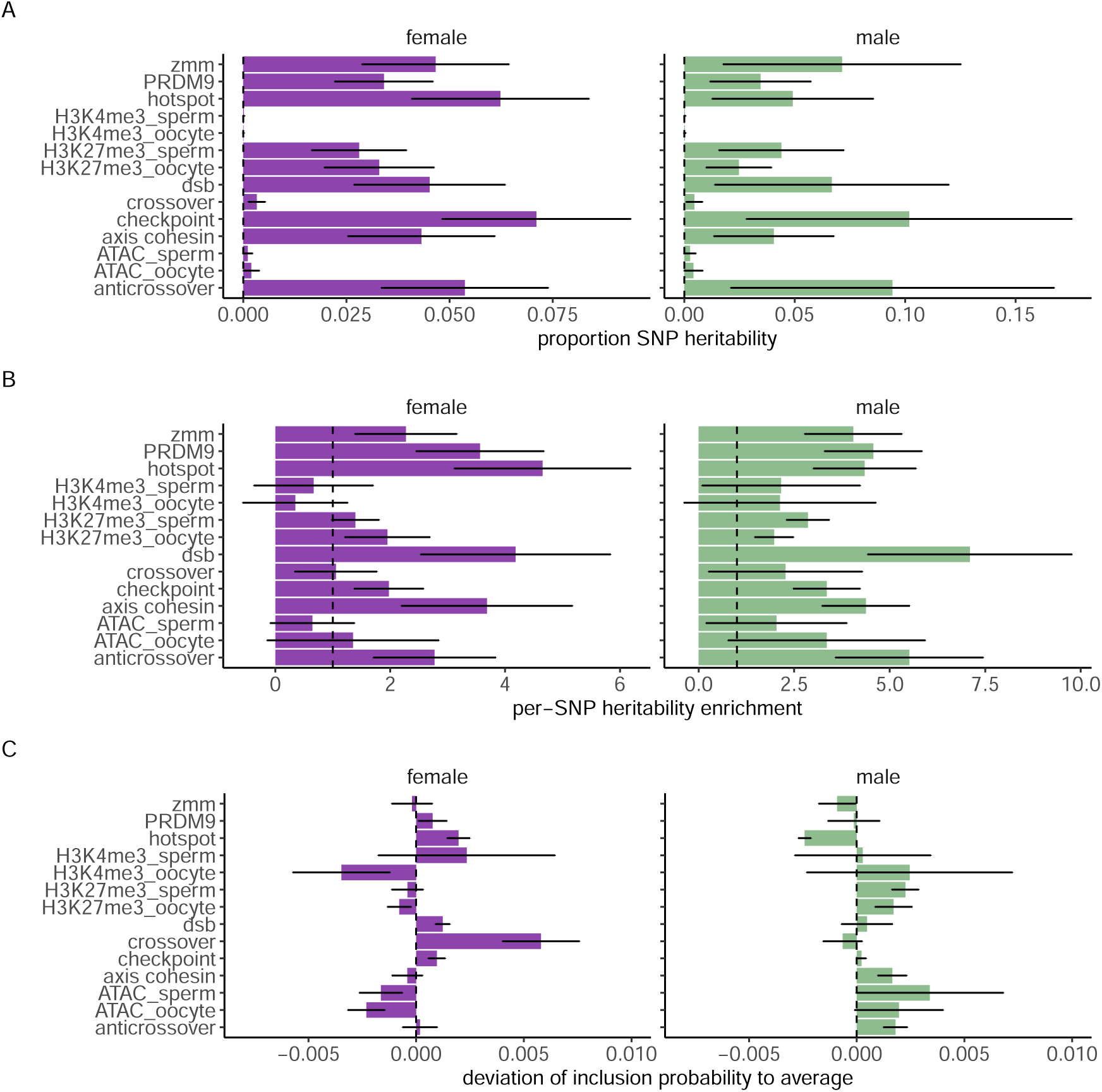
The proportion of variation in meiotic recombination rate attributable to gametic epigenetic marks and genes previously implicated in recombination. DNA loci are classified into their potential involvement in the following processes (see Table S11): histone methyltransferase gene (PRDM9); ZMM gene pathway crossover maturation (ZMM); hotspot localization (hotspot); double-strand break formation (dsb); crossover regulation designation (crossover); checkpoint and repair (checkpoint); Axis/Cohesin structure; regulation that is anti-crossover (anticrossover). Remaining labels refer to DNA loci in H3K4me3 and H3K27me3 epigenetic marks, as well as ATAC-seq ”tracks”, within both spermatocytes (pachytene stage) and oocytes. **(A)** From the meta-analysis estimates for recombination cross-over rate (CO), the proportion of phenotypic variation attributable to DNA loci for each annotation. **(B)** The per-SNP enrichment for each annotation group. **(C)** The deviation from the average inclusion of DNA loci in the model for each annotation group. Error bars give the SD intervals of the estimates. Estimates are made using SBayesRC, with full results given in the Data Table S1. More details on the statistical framework are given in the Materials and Methods.

In most eukaryotes, several recombination pathways that can result in crossover events operate in parallel during meiosis (*37*). Thus, local features may not be impactful on the overall scale of between-individual variation. We therefore also map SNPs to hypothesized global modulators of recombination. These pathways are hypothesized to have emerged independently in the course of evolution and perform separate functions, which directly translate into their roles in meiosis (*7*). We compile genes previously shown to encode proteins involved in meiotic recombination processes and use STRING gene interaction networks to map SNPs to their interacting neighbors. Genes are classified using their involvement in the following processes: double-strand break formation, axis/cohesin structure, hotspot localization and chromatin remodeling, ZMM gene pathway crossover maturation, anti-crossover regulation, crossover regulation designation, and checkpoint and repair (see Table S11 and Methods). Meiotic recombination double-strand breaks are also targeted by DNA sequence-specific binding of a meiosis-specific histone methyltransferase gene, *PRDM9*, which may drive broad-scale regulation of recombination in mammals (*38*). We also annotate SNPs to *PRDM9* binding sites defined using ChIP-seq peaks from HEK293T cells (see Methods). We find that all of these potential global modulators of meiotic recombination contribute a total of 37.7% (2.7% SD) and 47.1% (3.8% SD) to CO 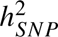 in females and males, respectively (Figure 4). This implies several known pathways operating in parallel to shape population-level variation during meiosis. Figure 4 also shows that enrichment is broadly similar across the sexes. We additionally examine the polygenicity for female CO, finding evidence that DNA variants on each chromosome contribute to variation in double-strand breaks that occur on other chromosomes (Figure S5). Taken together, this suggests that CO is a polygenic trait for which over half of the 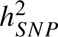 can be attributable to regions that have not been previously found to be implicated in the formation and modulation of double-strand breaks formed during meiosis.

## Polygenic prediction of meiotic recombination and relationships to other traits

We create a polygenic genetic predictor of female and male cross-over rates, which significantly predicts both female and male CO within an independent Scottish sample (GS), accounting for 2.1% and 1.1% of the phenotypic variance after correction for age and birth year, values which are in-line with expectations given the study sample size and the 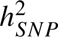 observed (Figure 5).

**Figure 5:**
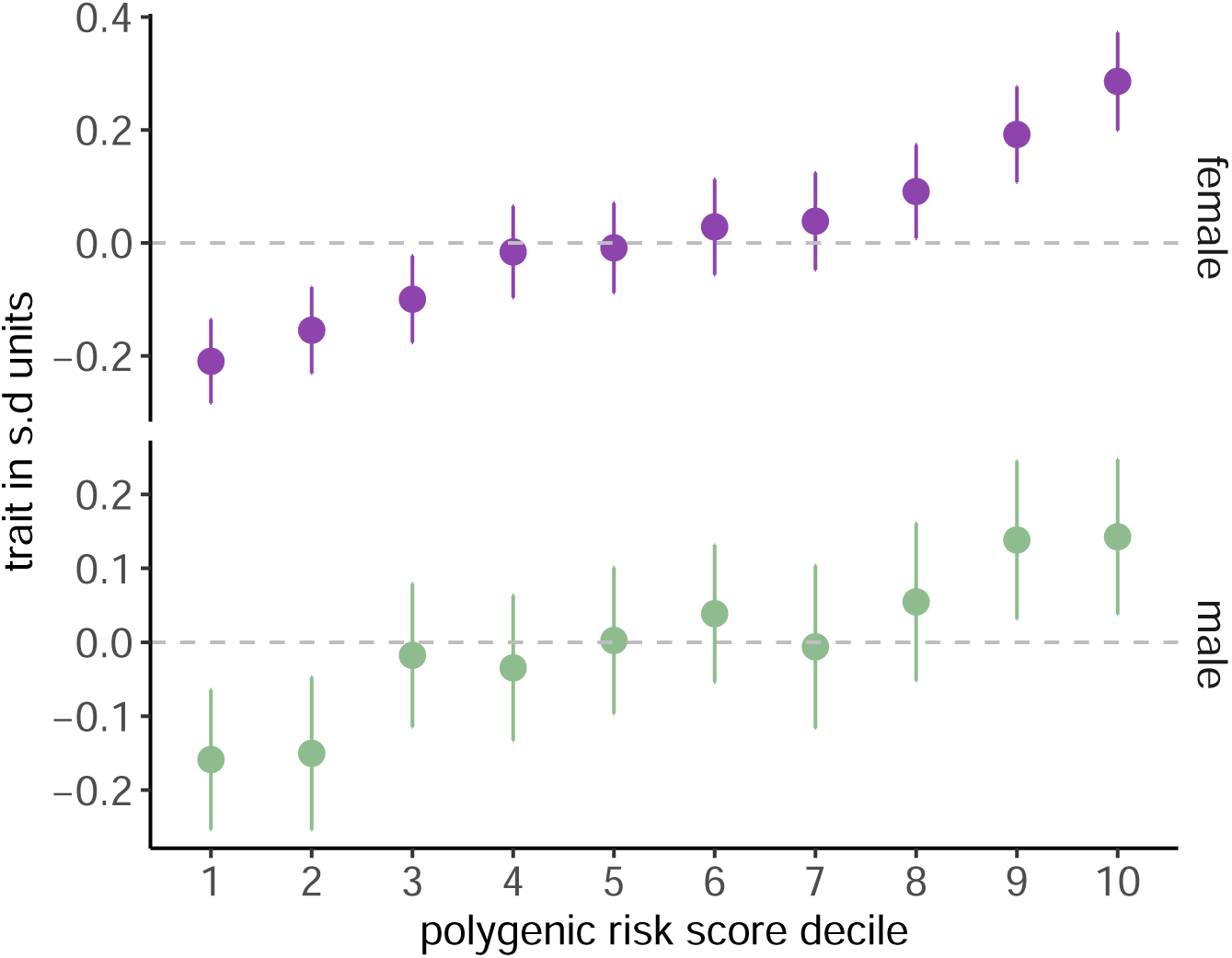
Polygenic prediction of meiotic recombination. For the measure of recombination cross-over rate (CO) a polygenic predictor significantly predicts trait value in a hold-out sample (a linear mixed effects model of CO for female slope = 0.996, SE = 0.114, 𝜒^2^ = 69.9, p-value < 2.2 × 10^−16^, male slope = 1.000, SE = 0.231, 𝜒^2^ = 18.7, p-value = 1.59 × 10^−5^). The mean trait value in SD units is given for each decile of the genetic predictor of CO for females in purple and males in green. Error bars give the 95% intervals of the estimates.

In the Supplementary Text, we further place our findings in the context of other studies, finding supporting evidence that the recombination count in an oocyte is correlated with maternal age (*10, 39, 40*) (Figure S6). We also examine potential relationships of female and male meiotic recombination and other fertility, mental and physical health, and behavioral phenotypes in UK Biobank and MoBa. Although we find associations with a range of traits, we caution that environmental, demographic and ascertainment factors may influence the links that we discover between recombination and other phenotypes, in addition to a potential underlying genetic relationship (see Figure S7 and Supplementary Text for a full discussion). Considering this, the most robust evidence for associations between meiotic recombination and other traits is between female CO and reproductive outcomes. After adjusting for a genetic predictor of study ascertainment, our genetic predictor of female CO is positively correlated with number of births in the UK Biobank, but negatively correlated with number of self-reported spontaneous miscarriages (UK Biobank code 3839). This is confirmed within MoBa by a positive phenotypic relationship between female CO and number of births, and a negative phenotypic correlation of CO with pregnancy complications as well as the time interval between pregnancies (Figure S7B). Our results are supportive of a hypothesis that because aneuploidy (incorrect chromosome number) in the fetus is one of the leading causes of pregnancy loss, women who produce eggs with more recombination may be expected to have more successful live births and show a reduction in pregnancy loss and pregnancy complications. Our findings also support a connection between female reproductive timing and DNA repair genes (*41*) (see Figure S7 and Supplementary Text for a full discussion). This suggests that a combination of genetic and environmental factors create relationships between meiotic recombination rate and female health and reproduction. The key next step is identification of the mechanisms that underlie these associations.

## Discussion

We present the largest meta-analysis of meiotic recombination to date, that is uniquely cross-cohort and utilizes a novel method to infer recombination events. We characterize the genetic architecture of recombination rate in males and females and discover seven novel associated DNA regions while replicating 29 out of 35 previously identified loci. Our findings confer that recombination is a polygenic quantitative trait, with genetic effects that differ among the biological sexes to some degree. Global modulators of meiotic recombination contribute relatively more than local features to 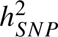, however, more than half of the 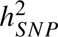 can be attributable to regions that have not previously been implicated in the modulation of recombination. Male meiotic recombination is slightly less heritable and potentially underlain by fewer regions of larger effect size. Thus, not only do recombinations yield genetic diversity, but they are also under genetic control and likely have had selective forces acting at the level of chromosome segregation. A cross-sex genetic correlation that is significantly less than unity, may permit independent selection responses and mechanisms to evolve that allow for the differences in meiotic recombination rates observed within the population. Finally, we created the first genetic predictor of recombination, allowing initial exploration into the relationship between recombination and reproductive health. Study limitations, such as the limitations of sample collection of both parents and children, are discussed in detail within the Supplementary Text. These will largely be addressed with whole genome sequencing data in multiple large cross-population samples of a range of genetic ancestry with improved assessment of potential ascertainment biases. Our study underscores how variation in recombination rates within individuals and across sexes impacts genome structure, disease risk, and reproduction in ways that we are only just beginning to understand.

## Acknowledgments

The Norwegian Mother, Father and Child Cohort Study is supported by the Norwegian Ministry of Health and Care Services and the Ministry of Education and Research. We are grateful to the families who take part in this on-going cohort study. We thank the Norwegian Institute of Public Health (NIPH) for generating high-quality genomic data. The research is part of the HARVEST collaboration, supported by the Research Council of Norway (#229624). We also thank the NORMENT Centre for providing genotype data, funded by the Research Council of Norway (#223273), South East Norway Health Authorities and Stiftelsen Kristian Gerhard Jebsen, and in collaboration with deCODE Genetics. We thank the Center for Diabetes Research, the University of Bergen for providing genotype data funded by the ERC AdG project SELECTionPREDISPOSED, Stiftelsen Kristian Gerhard Jebsen, Trond Mohn Foundation, the Research Council of Norway, the Novo Nordisk Foundation, the University of Bergen, and the Western Norway Health Authorities. The MoBa work was performed on the TSD (Tjeneste for Sensitive Data) facilities, owned by the University of Oslo, operated and developed by the TSD service group at the University of Oslo, IT Department (USIT). We thank Yunsung Lee for providing advice and R code for several of MoBa fertility phenotypes. Norwegian analyses were performed on resources provided by Sigma2 - the National Infrastructure for High-Performance Computing and Data Storage in Norway.

We thank and acknowledge the participants and investigators of the Generation Scotland Cohort study. Generation Scotland received core support from the Chief Scientist Office of the Scottish Government Health Directorates [CZD/16/6] and the Scottish Funding Council [HR03006]. Genotyping and methylation typing of the GS:SFHS samples was carried out by the Genetics Core Laboratory at the Wellcome Trust Clinical Research Facility, Edinburgh, Scotland and was funded by the Medical Research Council UK and the Wellcome Trust (Wellcome Trust Strategic Award “STratifying Resilience and Depression Longitudinally” (STRADL) Reference 104036/Z/14/Z). Analysis of the Generation Scotland data and the summary statistics obtained from the other analyses was conducted at IST Austria and is supported by the Scientific Service Units (SSU) of IST Austria through resources provided by Scientific Computing (SciComp).

We thank and acknowledge the participants and investigators of the Estonian Biobank (EstBB) study. The research was conducted using the Estonian Center of Genomics/Roadmap II funded by the Estonian Research Council (project number TT17). Estonian Data analysis was carried out in the High-Performance Computing Center cloud provided by University of Tartu.

The color scheme used throughout this paper of purple, green, and white were used as colors representing the suffragette movement.

## Author contributions

M.R.R. conceived and designed the study. I.K., L.H., G.P. and M.R.R conducted the analyses. R.J.H. developed the method to infer CO events. I.K. developed the computer code for the analysis, with input from M.R.R. P.K. provided the genomic annotations used within the paper. A.R., C.H., S.H., O.A., A.J., I.S., Y.L., A.D.A., Z.K., R.E.M., P.K., B.V.H., and A.H. provided study oversight and contributed to the data, analyses, and writing of the paper. Estonian data collection, genotyping and QC was conducted by the Estonian Biobank research team: Andres Metspalu, Lili Milani, Tõnu Esko, Reedik Mägi, Mari Nelis and Georgi Hudjashov. M.R.R., I.K., L.H. and A.H. wrote the paper. All authors approved the final manuscript prior to submission.

## Data and materials availability

Data from the Norwegian Mother, Father and Child Cohort Study is managed by the Norwegian Institute of Public Health. Access requires approval from the Regional Committees for Medical and Health Research Ethics (REC), compliance with GDPR, and data owner approval. Participant consent does not allow individual-level data storage in repositories or journals. Researchers seeking access for replication must apply via https://helsedata.no/. Information on how to access the MoBaPsychGen post-imputation QC data is available here: https://www.fhi.no/en/more/research-centres/psychgen/access-to-genetic-data-after-quality-control-by-the-mobapsychgen-pipeline-v/.

Estonian Biobank data (https://genomics.ut.ee/en/content/estonian-biobank) were used in this project. For access to be granted to the Estonian Biobank genotypic and corresponding phenotypic data, a preliminary application must be presented to the oversight committee, who must first approve the project. Ethics permission must then be obtained from the Estonian Committee on Bioethics and Human Research. Finally, a full project must be submitted and approved by the Estonian Biobank.

Access to the Generation Scotland data is available with appropriate permission from the Generation Scotland Access Committee. Applications should be made to. (https://genscot.ed.ac.uk/for-researchers/access).

Haplotype Reference Consortium Release 1.1 data (https://ega-archive.org/datasets/EGAD00001002729) are available by application to a Data Access Committee (DAC) of the Wellcome Trust Sanger Institute.

## Code availability

- THORIN software: https://github.com/RJHFMSTR/THORIN with user documentation: https://rjhfmstr.github.io/THORIN/.
- Custom R code to identify consensus double strand break events: https://github.com/RJHFMST R/Crossover_inference.
- Custom Bayesian multiple trait model code: https://github.com/medical-genomics-group/MAJA.
- Mobapsychgen pipeline: https://github.com/psychgen/MoBaPsychGen-QC-pipeline

## Materials and Methods

### The Norwegian Mother, Father, and Child cohort study: MoBa

The Norwegian Mother, Father and Child Cohort Study (MoBa) is a population-based pregnancy cohort study conducted by the Norwegian Institute of Public Health. Participants were recruited from all over Norway from 1999-2008. The invited women consented to participation in 41% of the pregnancies. The cohort includes approximately 114,500 children, 95,200 mothers and 75,200 fathers. The establishment of MoBa and initial data collection was based on a license from the Norwegian Data Protection Agency and approval from The Regional Committees for Medical and Health Research Ethics. The MoBa cohort is currently regulated by the Norwegian Health Registry Act. The current study was approved by The Regional Committees for Medical and Health Research Ethics (2016/1702).

Information from the Medical Birth Registry of Norway (MBRN), a national health registry containing information about all births in Norway, and the MoBa questionnaire data were used to identify sex, year of birth, multiple births (in the offspring generation), and reported parent-offspring relationships. Prior to genetic analyses, all pedigrees were constructed based on the reported relationships for each pregnancy. Wherever possible, biological sex was assigned using information from the MBRN. In instances where biological sex was not specified in MBRN, the reported biological sex from the MoBa questionnaires was used.

Behavioral and fertility phenotypes, such as smoking behaviors and pregnancy complications, were self-reported in the MoBa questionnaires or ascertained from MBRN (in the case of number of children). Smoking behavior and birth timing, or the time to pregnancy after partners first started trying, were reported in the weeks 15 and 30 of gestation questionnaires. Smoking during pregnancy was averaged across repeated measures in pregnancy while reporting ever smoking or smoking in the 3 months prior to pregnancy were only reported once. Mothers reported if they experienced any pregnancy complications in the 6-month questionnaire. Clinical diagnoses (eg. ADHD) were ascertained from the Norwegian Patient Registry (NPR) and measures of educational achievement and attainment from Statistics Norway (SSB).

Blood samples were collected from participating mothers and fathers at approximately the 17th week of pregnancy during the ultrasound examination. A second blood sample was taken from the mother soon after birth. The blood sample for the child was taken from the umbilical cord after birth. Biological samples were sent to the Norwegian Institute of Public Health where DNA was extracted by standard methods and stored (*42*). For more information on genotyping of the MoBa sample and for the family-based quality control pipeline used to prepare these data for analysis, see (*43*). Here, we use the data generated after completion of module0 through module5 of the MOBA psychgen genotype imputation QC pipeline (see Code Availability) but we replace the QC and imputation pipeline used in Ref. (*43*), with our custom pipeline for phasing and imputation as described below.

### The Estonian Biobank: EstBB

This project was granted ethics approval by the Estonian Committee on Bioethics and Human Research (https://genomics.ut.ee/en/content/estonian-Biobank). The activities of the Estonian Biobank are regulated by the Human Genes Research Act, which was adopted in 2000 specifically for the operations of the Estonian Biobank. Individual level data analysis in the Estonian Biobank was carried out under ethical approval 1.1-12/624 and 1.1-12/2856 from the Estonian Committee on Bioethics and Human Research (Estonian Ministry of Social Affairs), using data according to release application S16 from the Estonian Biobank.

As is common in many biobank studies, familial relationships were not known to us within the Estonian Biobank (EstBB). We calculated the coefficient of relatedness within the EstBB array data using the software KING (see Code availability). Parents were identified as individuals of opposite biological sex who: (i) were unrelated (relatedness IBS0 < 0.0012), (ii) were within 10 years of age, (iii) both share half of their DNA with the same individual; and (iv) who are both at least 15 years older than the individual that they share half their DNA with. The latter two criteria automatically selected the children. We are confident that 10,512 true trio families were identified this way and previous studies have also shown the accuracy of similar approaches to find trios (*44, 45*). We additionally identified a further 26,209 putative mother-child duos and 6,364 putative father-child duos, by selecting pairs of individuals that share half of their DNA and who are at least 15 years apart. This gave 96,682 individuals in total, which were genotyped with Illumina Global Screening (GSA) arrays.

### The Icelandic cohort: deCODE Genetics

The data from the Icelandic genealogy database were previously published (*10*). All biological samples used in this study were obtained according to protocols approved by the Data Protection Commission of Iceland and the National Bioethics Committee of Iceland. Informed consent was obtained from all participants, and all personal identifiers were encrypted with a code that is held by the Data Protection Commission of Iceland. In brief, 71,929 parent-offspring relationships (41,745 mother-offspring pairs and 30,184 father-offspring pairs) were available to estimate autosomal recombination across 690,421 phased autosomal SNPs that had been genotyped previously on a genome-wide Illumina BeadChip (HumanHap300, HumanHap300-Duo, HumanCNV370-Duo, Human610-Quad, Human1M or Human1M-Duo BeadChip). The double-strand break cross-over data presented in (*10*) we used to recalculate four phenotypes as described below.

### The Generation Scotland cohort: GS

Generation Scotland (GS) is a large population-based, family-structured cohort of over 24,000 individuals aged 18–99 years. The study baseline took place between 2006 and 2011 and included detailed cognitive, physical, and health questionnaires, along with sample donation for genetic and biomarker data. Ethical approval for the GS cohort was received from the NHS Tayside Committee on Medical Research Ethics (REC Reference Number: 05/S1401/89) and Research Tissue Bank status was granted by the East of Scotland Research Ethics Service (REC Reference Number: 20/ES/0021). Participants provided written informed consent.

Genotyping array data are available for 20,026 (83%) of the original GS participants. Samples were genotyped using the Illumina Human Omni Express Exome8V.1-2 A and Human Omni Express Exome-8V.1 A arrays. Quality control measures were implemented, filtering out samples with a call rate of < 98% and SNPs with a call rate of < 98%, HWE of < 1 × 10^−6^ and MAF of ≤ 1%, leaving 20,026 samples and 630,207 SNPs.

We calculated the coefficient of relatedness within the GS array data using the software KING. Parents were identified as individuals of opposite biological sex who: (i) were unrelated (relatedness < 0.0012), (ii) were within 10 years of age, (iii) both share half of their DNA with the same individual; and (iv) who are both at least 15 years older than the individual that they share half their DNA with. The latter two criteria automatically selected the children. We are confident that 2,680 true trio families were identified this way, which we used for our main analyses because we confirmed this using the available family information. We also identified an additional 3,273 mother-child duos and 1,185 father-child duos, by selecting pairs of individuals that share half of their DNA and who are at least 15 years apart and we also confirmed these relationships with the available family information.

### The UK Biobank

UK Biobank has approval from the North-West Multicenter Research Ethics Committee (MREC) to obtain and disseminate data and samples from the participants (https://www.ukbiobank.ac.uk/ethics/), and these ethical regulations cover the work in this study. Written informed consent was obtained from all participants.

We restrict our analysis to a sample of European-ancestry UK Biobank individuals to match the other cohorts analyzed. To infer ancestry, we use both self-reported ethnic background (UK Biobank field 21000-0), selecting coding 1, and genetic ethnicity (UK Biobank field 22006-0), selecting coding 1. We project the 488,377 genotyped participants onto the first two genotypic principal components (PC) calculated from 2,504 individuals of the 1,000 Genomes project. Using the obtained PC loadings, we then assign each participant to the closest 1,000 Genomes project population, selecting individuals with PC1 projection ≤ |4| and PC2 projection ≤ |3|. Samples were also excluded based on UK Biobank quality control procedures with individuals removed because of *(i)* extreme heterozygosity and missing genotype outliers; *(ii)* a genetically inferred gender that did not match the self-reported gender; *(iii)* putative sex chromosome aneuploidy; *(iv)* exclusion from kinship inference; *(v)* withdrawn consent.

We use genotype probabilities from version 3 of the imputed autosomal genotype data provided by the UK Biobank to hard-call the single nucleotide polymorphism (SNP) genotypes for variants with an imputation quality score above 0.3. The hard-call-threshold is 0.1, setting the genotypes with probability ≤ 0.9 as missing. From the good quality markers (with missingness less than 5% and 𝑝-value for the Hardy-Weinberg test larger than 10^−6^, as determined in the set of unrelated Europeans) we select those with rs identifier, in the set of European-ancestry participants, providing a dataset of 23,609,048 SNPs.

### Phasing and haploid imputation within families

For MoBa, EstBB and GS, we performed inter-chromosomal phasing of the family genetic data using SHAPEIT5 (*45*) (see Code availability). Since we have access to parental genomes, we perform what we refer to as ”pedigree-based phasing”, meaning that we explicitly estimate offspring haplotypes from their parents. This requires a three-column pedigree file listing offspring, fathers, and mothers. In cases where one parent is unavailable, the missing parent is indicated by ”NA”. This procedure results in phased genotype data where the paternally and maternally inherited haplotypes are inferred. The key advantage of using pedigree-based statistical phasing over traditional Mendelian logic is its ability to infer the parent-of-origin (PofO) of alleles even when both parents are heterozygous, which is accomplished by leveraging haplotype information from the broader population, providing more robust and accurate results, such as done in traditional statistical phasing.

For MoBa, we used only trios where parents and children were genotyped on the same genotyping array and conducted the phasing separately for each array in six sets. For EstBB and GS, as we wished to maximize both the number of trios and sibling pairs, we phased both trios and duos that were genotyped on the same array. Thus phasing was conducted within-family and within-array, within each study.

We then matched the phased data to Genome Reference Consortium Human Build 37 (GRCh37) and swapped reference alleles to the Haplotype Reference Consortium data EGA release (*46*) (see Data and materials availability). We then imputed SNPs in the parents and the children using IMPUTE5 v1.1.4 with the Haplotype Reference Consortium as a reference panel. As our data is phased, with each haplotype assigned to a specific parent, we used the parameter -out-ap-field to run a haploid imputation of the data and separately imputed the paternal haplotype and the maternal haplotype as done previously (*12*). As a result of haploid imputation, the PofO of imputed alleles can be probabilistically deduced from the imputation dosages: an allele imputed with a dosage of 0.9 on the paternal haplotype has 90% probability of being inherited from the father (i.e., PofO probability = 90%). This was done in 448 segments of the DNA for each set of data, and thus was also within-array. For each segment, we removed variants with an INFO score < 0.9, minor allele frequency < 0.1%, keeping G/T SNPs only after imputation. Within each study, we concatenated files across segments for each chromosome. For MoBa, we then additionally merged the files across the six genotyping array sets for each chromosome. A final round of QC was conducted again, keeping only SNPs with missingness < 5%, INFO score > 0.9, minor allele frequency > 1% and HWE p-value > 5x10^−8^. If there were multiple genotyping arrays within a biobank as in the case for MoBa and EstBB, we calculated fixation index (FST) across the array sets and excluded SNPs that show any signs of being differentiated with FST value > 0.001. We also calculated the heterozygosity of each individual genome-wide using the plink –het command, and excluded individuals based on the F-statistics of heterozygosity.

For full methodological details on phasing and imputation in Iceland, see Ref. (*10*).

### Identification of meiotic crossovers

For MoBa, EstBB and GS, we infer crossover positions using Identity-By-Descent (IBD) mapping. Identity-by-descent (IBD) mapping is a powerful approach for identifying haplotype segments co-inherited from a common ancestor among pairs of relatives. To do this, we used the THORIN v1.2 tool (*12*) to map haplotype segments shared IBD. As we performed pedigree phasing of our data, the offspring are perfectly phased and thus we observe no parental recombination in the offspring haplotypes. Hence, we map IBD in the parents to identify crossovers, using the offspring as reference individual. By analyzing haplotype segments shared IBD with the same set of parents across the 22 autosomes for a given offspring, we can determine which haplotype segments are inherited from the same parent. This enables the construction of parental haplotype sets, which can then be used to perform inter-chromosomal phasing and identify the segregating haplotypes inherited from each parent across the genome. This approach goes beyond traditional phasing methods, which are limited to resolving haplotypes within individual chromosomes (intra-chromosomal phasing).

Identifying the IBD segments then allows us to infer crossover positions in the gametes transmitted from each parent to their offspring. For each chromosome and each meiosis, THORIN gives the per-variant site IBD sharing probability and IBD segments. We then use a custom approach (see Code Availability) to identify consensus double strand break events, by identifying a break in IBD segments occurring simultaneously on both haplotypes where the IBD segment ends on one haplotype and starts on the other haplotype. This outputs the start and end of crossover positions and is inspired from Ref. (*12*) with specific modification. We identify consensus crossovers, which are those that are inferred on both the paternal and maternal haplotype with a distance smaller than a threshold of 3cM. We exclude segments of IBD that are less than 3cM in length and we only consider segments that are more than 3cM apart as being independent events. Thus, we assumed that all crossovers were Class I crossovers and subject to crossover interference and that short crossovers (those less than 3cM) were indicative of non-crossover gene-conversion events, non-interfering Class-II crossovers, phasing errors, and/or genotyping errors. This minimizes the likelihood of switch errors in the parents that would result in false positive detections (*12*).

For full methodological details on cross-over inference in Iceland, see Ref. (*10*).

### Recombination rate phenotypes

Here, we use the obtained datasets obtained within each population to calculate four phenotypes of female and male recombination:

1. **CO**: the total number of double-strand break crossovers occurring on the autosomes for each meiosis.
2. **PA**: the proportion of the gamete’s paternally or maternally derived genome that is from a single grand-parental origin. This represents a measure of the intra-chromosomal allelic shuffling, which is defined as the probability that two randomly chosen loci on the same chromosome are uncoupled in meiosis. As crossover positions are determined for individual gametes, and the parental origins of the chromosome segments delimited by these crossovers are known, then the proportion of the gamete’s genome that is from a single grand-parental origin can be determined. This proportion is a measure of the probability that the alleles of a randomly chosen pair of loci are shuffled during formation of the gamete (equivalently, the proportion of locus pairs at which the alleles are shuffled), which is referred to in the literature as 𝑟_𝑖𝑛𝑡𝑟𝑎_ (*13*). In practice, we sum the cM lengths of the segments of each chromosome that are attributable to each grandparent. We then calculate the fraction of the gamete’s genome at each chromosome that is from a single grand-parental origin. For example if the chromosome is 70% grand-paternal and 30% grand-maternal, then the value 0.7 is used, representing the percentage that is of a single origin. For each meiosis, the average is taken across chromosomes.
3. **MD**: The distance (in bp) between consecutive crossover events which is used as a proxy for interference. Distances are measured in bp between consecutive events on each chromosome. For each meiosis, the average is taken across chromosomes.
4. **TD**: The average relative distance of the crossovers from the telomere. Distances are measured in cM to the closest telomere on each chromosome arm. Previous work (*7*) in other species has taken the minimum from both arms of the metacentric chromosomes and the minimum from acrocentric chromosomes (so whichever arm had the smaller distance) and here, we follow the same approach. For each meiosis, the values are then averaged across all arms.

For deCODE, the same four phenotypes were constructed using the data from Ref. (*10*).

### Association testing

We conducted genome-wide association testing for each of the four phenotypes and each sex separately. For deCODE data, the analysis pipeline described in Ref. (*10*) was used, with summary statistic data obtained using the mixed-linear model association testing software BoltLMM. For MoBa and EstBB, custom mixed-linear model association testing software was used, which matches Regenie, but with the leave-one-chromosome-out genomic predictor obtained from the first step of Regenie being replaced by those obtained from a Bayesian analysis, as described below. Summary statistics across the three studies were meta-analyzed with fixed-effect inverse variance weighting, using the software METAL. The 𝜒^2^ statistics for the meta-analysis divided by their expectation is given in Figure S2 and the median and mean value for each study is given in Table S9.

To these summary statistic data, we apply the GCTA cojo algorithm of Ref. (*47*) to perform a stepwise model selection procedure to select independently associated SNPs. We use the default threshold p-value of 5 · 10^−8^ to declare genome-wide significance, the default distance of 10 Mb for the complete linkage equilibrium assumed between SNPs and default collinearity multiple regression 𝑅^2^ between the SNPs that have already been selected and a SNP to be tested of 0.9. The DNA regions identified are presented in Table 1 for each recombination measure, the number of SNPs with genome-wide significant marginal test statistic are given in Table S2 and the GCTA-cojo selected SNPs within each region are given in Table S3.

### Between-sex genetic correlations

To estimate the genetic correlations between the sexes, we use a multi-trait Bayesian sampler, MAJA (*33*). The Gibbs sampler, an iterative Markov Chain Monte Carlo method, is based on a linear regression model. For each meiotic recombination measure, the phenotypes of men, 𝒀_𝒎_, and women, 𝒀_𝒘_, are stacked so that 𝒀 is of dimension (𝑛_𝑚_ + 𝑛_𝑤_) × 1, where 𝑛_𝑚_ and 𝑛_𝑤_ refer to the number of male and female individuals. The phenotypes are linearly related to the genotypes of men, 𝑿_𝒎_, and women 𝑿_𝒘_:

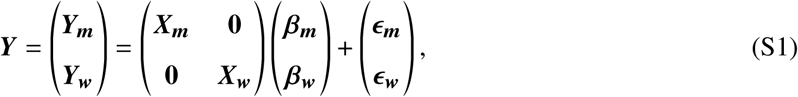

The effects of each marker 𝑗 on men and women, 𝜷_𝒋_ , are modeled using a multivariate spike-and-slab prior distribution to accommodate zero effects sizes

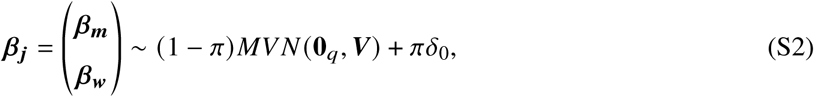

where 𝜋 is the marker exclusion probability common to all phenotypes, 𝑀𝑉 𝑁 (**0**_𝑞_, 𝑽) is a multivariate normal distribution with mean 0 and variance-covariance 𝑽 and 𝛿_0_ the Dirac delta distribution. A marker is not included in the model if the effects of all traits are estimated to be 0. Through sampling the effects of each marker conditional on all the other markers, correlations between the markers (linkage disequilibrium) are automatically taken into account. The variance-covariance of effects, 𝑽, and the variance-covariance matrix of 𝝐 are modeled as outlined in Section 2 of Ref. (*48*), using a modified Cholesky decomposition. The variance-covariance of effects, 𝑽, models the variances of the sexes as well as the covariance between the sexes. The correlation is then calculated from the estimated variances and covariances, and their standard deviations. The sampler assumes that the residual error is independent between the sexes which is reflected in the prior Gamma distribution of the variances of 𝝐.

We apply the model to SNPs with minor allele frequency > 5% in MoBa and > 10% in EstBB, where access to individual-level data was obtained and sample sizes are sufficient. Posterior mean estimated effect sizes are used to create the leave-one-chromosome-out predictors that are used for the association testing.

### Cross-trait genetic correlations

We also use MAJA to estimate the genetic correlations between the four traits within each biological sex as:

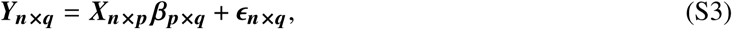

where 𝒀_𝒏_**_×_**_𝒒_ is a matrix of 𝑞 phenotypes for 𝑛 individuals, 𝑿_𝒏_**_×_**_𝒑_ is the genotype matrix for 𝑝 markers, 𝜷_𝑝×𝑞_ denotes the effect sizes for 𝑞 traits and 𝝐_𝒏_**_×_**_𝒒_ represents the residual error matrix. Each column of the 𝒀_𝒏_**_×_**_𝒒_ and 𝑿_𝑛× 𝑝_ matrices is standardized. The effects of each marker 𝑗 on the multiple phenotypes, 𝜷_𝒋_ , are modeled as the right side of Equation S2. The model also determines the variance-covariance of the residual errors across traits.

### Heritability and genetic correlations with SumHer

We estimate the proportion of phenotypic variance attributable to the SNPs (the SNP heritability, ℎ^2^ ) and the genetic correlations within and between women and men using the SumHer software (*49*). We apply the model to the summary statistic data from within each cohort and to the meta-analyzed estimates. We use a subset of 6 million common SNPs (frequency 1% and above) that are shared across cohorts and the UK Biobank.

### Polygenicity of female recombination rate with GMRM

Female CO was calculated in the Estonian Biobank (EstBB) 22 times, each time for all chromosomes other than a focal chromosome. We then ran 22 models, regressing each CO measure onto two groups of covariates jointly, the genotypes of the focal chromosome and the genotypes of all other chromosomes. For example for chromosome 1, the double strand breaks occurring on chromosomes 2-22 were counted and regressed onto two groups of covariates jointly, the genotypes of chromosome 1 and the genotypes of chromosomes 2-22. Thus, we estimate the degree to which SNPs on a specific chromosome contribute to the variation in CO observed in other chromosomes. The model was run using the GMRM software (*50*) on the same set of SNPs used in MAJA as described above.

### Heritability partitioning and genomic prediction with SBayesRC

We estimate the proportion of phenotypic variance attributable to the SNPs (the SNP heritability, 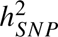) and partition it across genomic annotations using the SBayesRC software (*34*). We use a subset of 6 million common SNPs (frequency 1% and above) that are shared across cohorts and the UK Biobank.

Genomic annotations were constructed using the hg19 assembly. We make a list of genes known to encode proteins involved in meiotic recombination or closely related processes (Supplementary Table S11). Polygenic pathway annotations were generated by selecting seed genes encoding known components of seven key meiotic processes (DSB formation, axis/cohesin structure, hotspot localization, ZMM crossover maturation, anticrossover regulation, crossover regulation designation, and checkpoint/repair), expanding these sets to include direct and secondary interactors using the STRING v12 database (*51*), and defining the final genomic features as gene bodies padded by 100 kb upstream and downstream.

Histone modification (H3K4me3, H3K27me3) and chromatin accessibility (ATAC-seq) data for sperm were obtained from GSE68507 (*52*) and GSE120507 (*53*), while corresponding data for germinal vesicle stage oocytes were obtained from GSE124718 (*54*), and PRDM9 binding sites were defined using ChIP-seq peaks from HEK293T cells (GSE99407) (*55*). To ensure unbiased comparisons of heritability enrichment, epigenetic tracks were calibrated by first expanding narrow peaks (H3K4me3, ATAC, PRDM9) by 500 bp and broad peaks (H3K27me3) by 1000 bp, removing regions overlapping the ENCODE hg19 blacklist, and subsequently downsampling the sex with higher coverage to match the total base-pair footprint (estimated using 1000 Genomes Phase 3 EUR variants (*56*) of the limiting sex via a binary search of top-ranking peaks. We calibrate these annotations in terms of base pair coverage between male and female sides and then map the SNPs to these.

We fit all of our custom annotations jointly, alongside a subset of the annotations proposed in the SBayesRC paper (*34*) including all those from UCSC (coding, intronic, promoter, 3’ and 5’ UTR, synonymous, non-synonymous), ENCODE transcription factor binding sites, as well as ten bins of minor allele frequency.

We use the posterior mean SNP effects obtained from SBayesRC to predict female and male CO within the GS data, with a mixed-effect linear model fitting a random effect term for parental identity to account for repeated measures, as well as fixed effect terms for parental birth year and parental age, implemented using the R package lme4. We employ the implemented anova function to test for significance (𝜒^2^ probability) of a model parameter against a null hypothesis, which does not include the model parameter.

We also predict CO in the UK Biobank data and then estimate the correlation between female and male meiotic recombination and a wide-range of reproductive outcomes (full trait codes and estimates are given in Supplementary Data). UK Biobank phenotypes are adjusted by sex, birth year, genotype batch, north-south and east-west residential location and 20 PCs of the genomic data. Additionally, we create a genetic predictor of study ascertainment, using the SNP effect estimates of Ref. (*57*) and include this within the model as an adjustment variable. SNP associations and SNP heritability estimates are likely less impacted by ascertainment weighting, but substantial discrepancies are likely when studying correlates of meiotic recombination with behavior, lifestyles and social outcomes (*57*) (see Supplementary Text).

### Testing for sex-specific SNP effects

To test for SNP-by-sex interactions, we use two methods:

1. A z-score approach: 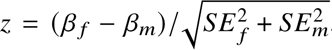, with 𝛽 the meta-analysis regression coefficient of females (f) and males (m) and 𝑆𝐸 the standard error. This assesses the difference between the meta-analysis regression coefficients, assuming a normal distribution. We conduct this test for each of the four traits.
2. Within-study interaction testing using a mixed-effect linear model implemented using the R package lme4. We fit a term for sex, each SNP, sex·SNP interaction, and one for parental identity to account for repeated measures. We employ the implemented anova function to test for significance (𝜒^2^ probability) of the interaction term against a null hypothesis, which does not include this term.

The results of this testing are presented in Table 3.

### Fine-mapping using SuSiEx

We perform fine-mapping across men and women by re-purposing SuSiEx (*14*) using meta-analyzed effects and the linkage disequilibrium matrix for men and women in the UK Biobank. The bp regions for fine-mapping are selected based on the lead associated SNP identified per region across all traits and sexes ±5 Mb (± 10 Mb for chromosome 6 and 17). The causal variants are selected based on 95% confidence sets. The sex-specific causal probability is set to be greater than 0.9 and SNP identities of high probability are given in Table S4.

## Supplementary Text

### Different ways of calculating recombination phenotypes

To investigate whether our results are sensitive to the method of calculation, we take advantage of recently curated sex-specific recombination maps in Iceland (*2*) and the most recent association study data from Iceland (*10*) to compare the genetic effect estimates of recombination phenotypes calculated in different ways. Irrespective of whether the proportion of single ancestry transferred is calculated as we do here, or by dividing the estimate of each chromosome by the length as suggested in the literature (*13*), the genetic estimates are not distinguishable from being identical for females (SumHer estimated genetic correlation = 0.998, 0.033 SE) or males (SumHer estimated genetic correlation = 1.021, 0.034 SE). Likewise, the genetic effects for distance from the telomere (TD, SumHer estimated genetic correlation = 0.988, 0.033 SE for females and 0.898, 0.087 SE for males) and mean distance among consecutive recombination events (MD, SumHer estimated genetic correlation = 0.986, 0.039 SE for females and 0.973, 0.034 SE for males) are also indistinguishable, whether they are calculated in cM or base pairs.

We select the same two measures of CO and TD used in the most recent genetic association study of recombination rate (*10*) and combine this with two additional measures that have not yet been used in large-scale human association studies: the PA measure (*13*) and the MD measure. The PA measure has also been used across studies of non-human organisms (*7*). We use MD as a measure of interference. We do not calculate a recombination hotspot phenotype used in Ref. (*10*), which describes differences among individuals in the likelihood that cross-overs occur within sex-specific hotspot locations, or the GC content and ENCODE replication timing score measures used (*10*). Sex-specific recombination maps with whole genome sequence data are not available outside of Iceland and we wished to avoid the complication of standardizing hotspots, gc content and ENCODE scores measures across studies. As we do not have sequence data and genomic annotation for all studies, we felt cross-study comparisons for these location-based measures are best left to future studies when such data are available.

### Limitations in the representative nature of our findings

We use a Hidden Markov Model (HMM) method, to conduct identity-by-descent mapping. This requires only a duo (parent–child pair) to call crossovers, but will have greatest power for duos from informative nuclear families or families in which the grandparents or other more distant relatives are also genotyped (*8, 10, 58*). Previous work has shown the effect on the genome-wide recombination rate is negligible for association testing (*8, 10*). Like other studies, we minimize this effect by removing double crossovers called within short intervals (see Methods), which are likely switch or genotype errors. This has been shown to homogenize estimates across cohorts (*9,40*) and we find the same here. We conservatively focus on reliably called double-strand breaks that can be studied across cohorts so that association study testing power can be improved. However, as a result, our estimates will likely be an underestimate: we overlook complex cross-overs and gene-conversion events that were previously studied (*10*). In particular, complex cross-overs may be those which increase in frequency in later-life (*10*) and this may be why observed increases in CO with age are reduced as compared to other studies (see below).

Neither our results, nor those of other studies (*8–10,40*), directly observe recombination rates in sperm and oocytes, because they all measure recombination phenotypes in live births. This is a form of ascertainment bias, where our observations only capture variation in double strand breaks that can be detected from genotyping arrays, in live births only for families where multiple individuals consent to join a biobank. Thus, the variation we observe is likely only a small fraction of the true underlying variation in meiotic recombination, as it is the variation observed after the selective process of a gamete becoming a live birth and the family enrolling for study. If fetal survival depends upon the amount of recombination, as it likely does, the observed effects on numbers of crossovers may not reflect changes in the recombination rate in oocytes. It could be possible instead to observe recombination in sperm and oocytes directly, but sampling and study enrollment would likely be difficult. To date this has only been conducted using individuals attending fertility clinics (*16*), which is another biased sample containing many individuals with fertility problems.

Nonetheless, our work is still of inherent biological interest because it reflects the variation observed in the population among those that are born, which as we have shown here is likely polygenic, attributable largely to regions that have been experimentally shown to be involved in meiotic recombination (Figure 4) and is potentially related to other outcomes (see below for further discussion).

### Differences across cohorts

In this study, we attempt to eliminate as many confounders as we can by applying the same methodology, as far as possible, across cohorts, and by focusing our analyses on only three large cohorts, each with tens of thousands of families. In the main text, we present evidence that the distributions of the recombination phenotypes that we calculate (Figure 1 and Table S1) have similar means and variances across cohorts. We show that the variation we see among individuals within the three cohorts, is attributable to common SNPs in the same proportion (Figure 3A), with similar estimated between-sex genetic correlations (Figure 3B) and between-trait phenotypic and genetic correlations (Figure S1). Nevertheless, although the individual-level variance appears to be similar, there are slight differences in population-level means with age (Figure S6). There could be numerous reasons why we observe these mean cohort differences, which warrant further discussion.

We can mostly exclude differences in crossover-calling methodology or in statistical approaches, but study heterogeneity may be attributable to hidden confounders, including, but in no way limited to differences in: family data ascertainment, assisted reproductive technologies, oral contraception, pregnancy terminations, trends through time, or simply to chance effects in relatively small samples. Studies differ in the degree of relatedness, with MoBa consisting of trios, where as the EstBB, GS and Icelandic datasets consist of a mixture of extended relationships, trios (2,600 in GS and 10,500 in EstBB) and many thousands of duos. MoBa is the only data where both parents are always observed, but the percentage of repeated birth observations is lower than the other cohorts and there are far fewer births below the age of 24 (Figure S7). We observe many more mother-child duos with repeated maternal observations, than for father-child duos in the other cohorts (Table S1). Thus, family structure differs across cohorts, which likely induces estimation error which may vary with age.

There are also inter-cohort differences in reproductive timing and in family size, which has previously been found to be associated with recombination rate. For example, if women with higher baseline recombination rates have more children later in life, the number of recombinations will appear to be higher in older mothers. There is some evidence that previous analyses could be confounded by this, if analyses do not account for the baseline rate of the mother (*40*). For example, in consumer genomic testing data, age relationships can disappear in data where multiple pregnancies are observed, but it is unclear if this is an effect of reduced power (*9*). Previous work has shown that a maternal age effect on the number of crossovers is small and positive, and that the differences between cohorts are likely due to chance effects in small samples (*9*). Therefore, we use linear mixed effect modeling throughout the paper, where relationships and age effects are always considered, whilst accounting for the repeated measures and intra-individual variation. We could not explicitly include family size as a covariate in our analyses, since we did not have that information, but we should be capturing this effect by including the mother/father effect in our mixed effect model. Since the estimate for each mother is drawn from the same distribution as the other mothers in the cohort, we might expect some shrinkage towards the mean mother effect. This could potentially, for example, reduce the estimate for high-recombining mothers and thus inflate the fixed effect estimates, but we do not think this drives any of the relationships we present here.

Oral contraceptive use, the choice of reproductive timing across cohorts, the degree of study ascertainment and the availability of healthcare may all differ across cohorts. The decades in which the sampling periods occurred differ across studies and individuals may make different reproductive choices, the availability of pregnancy terminations, the introduction of prenatal screening, and general healthcare practices will change through time. All of these factors can contribute to differences among cohorts in the relationship of recombination rate to parental age. In particular, many forms of contraception suppresses ovulation. So, if eggs are released in the order they are produced during fetal development, as happens in mice, the age of eggs released at a certain point in a woman’s lifetime will differ depending on whether and for how long she has taken oral contraception. This could confound the measured effect of maternal age on the number of crossovers, but evidence for this is mixed at present (*59*). We could not control for contraceptive use as we did not have this information within our data, but we did adjust for birth year of the parent.

Finally, the number of crossovers is more over-dispersed in females than in males, relative to the mean. We should thus expect more variation, and less precise estimation, in females than in males. Thus, differences we see between cohorts may simply reflect chance effects.

### Relationship of recombination with maternal age and fertility phenotypes

In humans, there is evidence that the recombination count in an oocyte is correlated with maternal age (*10, 39, 40*). We also find evidence of age effects on the female recombination rate in EstBB (P = 1.596 · 10^−11^) and MoBa (age^2^ P = 0.001944), but not in GS. Overall, across populations we find lower recombination in live births of young mothers and highly variable recombination rates in the live births of older mothers across cohorts (Figure S6). This is supported by previous studies showing high heterogeneity across cohorts in the number of cross-overs observed in later-age births (*40*), which likely reflects a wide variety of differences across studies in reproductive timing and study ascertainment as discussed above. However, a general pattern of increased recombination rates in the live births of older mothers is evident and supported by evidence that maternal age impacts mean distance between consecutive crossovers negatively (P_EstBB_ = 2.642 · 10^−5^, P_MoBa_ = 0.003697). This also supports the finding in Ref. (*9*) that there is an increasing number of crossover events escaping interference, thus decreasing the distances between the events, with maternal age. This correlation is thought to result, not from the recombination rate of eggs increasing with age, but rather from a high recombination count increasing the chance of a gamete becoming a live birth, which becomes more pronounced with advancing maternal age (*10,39,40*). A higher recombination frequency along a chromosome may reduce the chance of maternal age–related non-disjunction, as absent or reduced levels of recombination, along with sub-optimally placed recombinant events, increase the likelihood of mal-segregation (*60*).

Study ascertainment biases in genome-wide association studies have been shown to influence estimated relationships among phenotypes, especially behavior, lifestyles and social outcomes (*57*). Here, we take care to correct as best we can for sample ascertainment in the UK Biobank when examining relationships of our genetic predictor of meiotic recombination and other outcomes (see Figure S7 and Methods). We create a genetic predictor of study ascertainment, using the SNP effect estimates of Ref. (*57*) and include this within the model as an adjustment variable. Our aim is to reduce the likelihood of discovering trait correlations that are driven by similar forms of study ascertainment across cohorts (say of healthier individuals, with higher socio-economic status). For example, when we do not include the genetic predictor of study ascertainment then our predictor of female CO is strongly associated with educational attainment (𝑝 = 4.87 × 10^−09^), which then disappears entirely when including the adjustment variable (Figure S7A).

Women who produce eggs with more recombination may be expected to have more successful live births and show a reduction in pregnancy loss and pregnancy complications since aneuploidy (incorrect chromosome number) in the fetus is one of the leading causes of pregnancy loss. We find that our genetic predictor of CO is positively correlated with number of births in the UK Biobank (Figure S7B). This is confirmed by a positive phenotypic relationship within MoBa. Our genetic predictor of CO is also associated with a lower number of self-reported spontaneous miscarriages in the UK Biobank data (UK Biobank code 3839). Supporting this, we find a negative phenotypic correlation of CO and pregnancy complications, and CO with birth intervals, within the Norwegian data (Figure S7B).

Recombination rates are expected to be influenced by environmental and demographic factors and may be linked to other phenotypes (*7*). In the UK Biobank, we find our genetic predictor of female CO also correlates with height, bone mineral density, lipid levels and smoking prior to pregnancy (Figure S7A). The polygenic predictor of male CO is also associated with bone mineral density (Figure S7A), supporting a potential link between fertility and osteoarthritis (*61, 62*). Previous studies have shown that reproductive timing and premature ovarian insufficiency are associated with several traits such as cardiovascular disease, dyslipidemia, and osteoporosis (*63*). Our findings also support a connection between menopause and DNA repair genes (*41*), where women who produce eggs with more recombination may be expected to have more successful live births and a shorter reproductive period (Figure S7). Recent findings (*17*), actually show that relationships between female reproductive traits, aneuploidy and reproductive timing differ across genes (see Figure 4 of Ref. (*17*) for example, where the pattern at *SMC1B* differs to that of all other genes). Our estimates reflect the phenotypic relationships that are expected when trait correlations are inferred genome-wide.

Thus, our findings speculatively suggest an overlap of female meiotic recombination, aneuploidy, and reproductive aging, that is linked to pregnancy loss within our species. However, given the strong sensitivity of the results to correction for ascertainment, we caution that these are all observational-level correlations and confounding factors cannot be ruled out. These data only suggest that a combination of genetic and environmental factors create relationships between meiotic recombination rate and female health and re-production but future studies are needed that do more to explore potential ascertainment and confounding factors and that focus on the potential mechanisms involved.

## A Supplementary Figures

**Figure S1:**
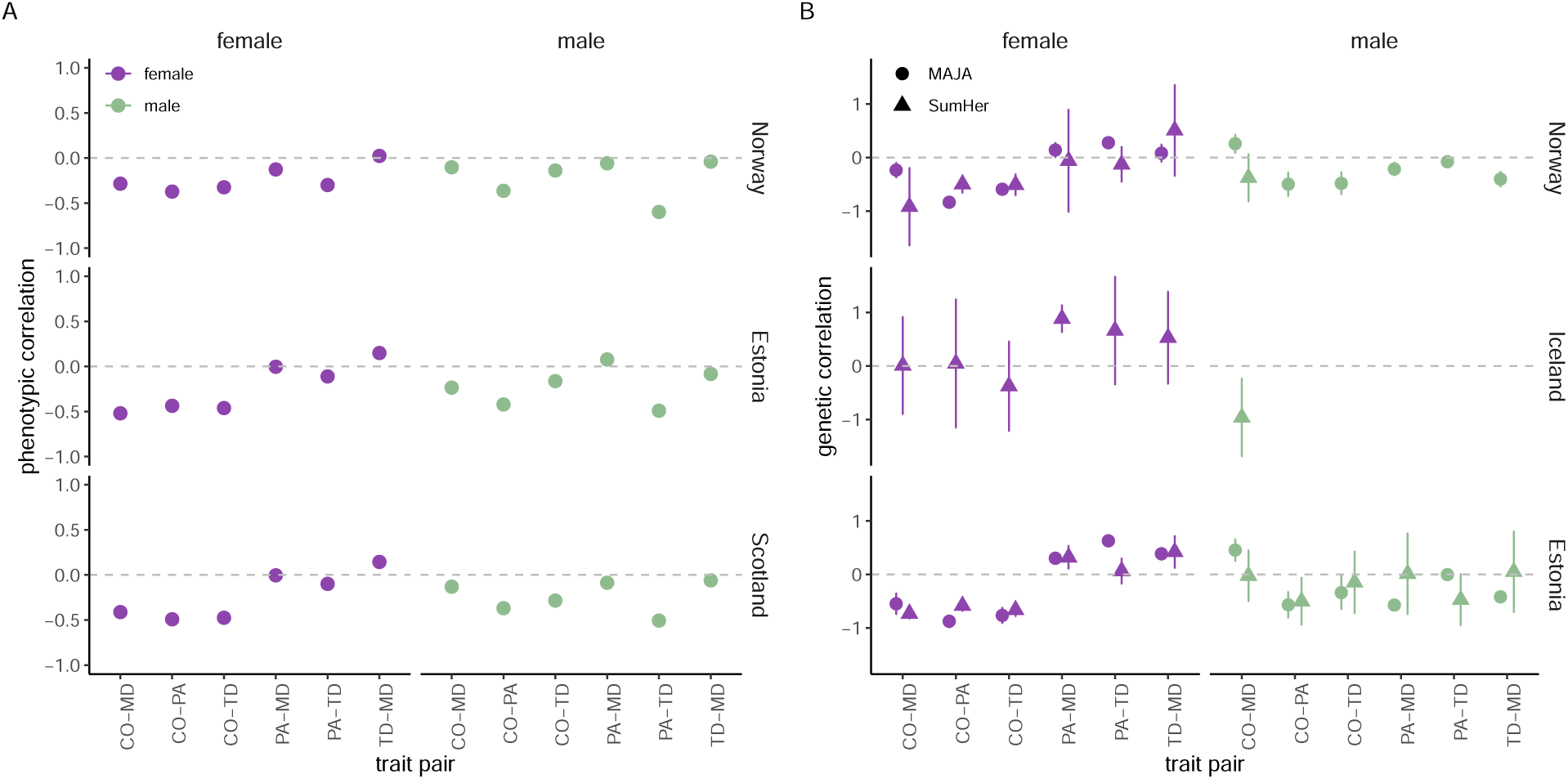
Phenotypic and genetic relationships among four recombination phenotypes across countries. In (**A**), phenotypic correlations among four meiotic recombination phenotypes across three countries, within individuals of female and male biological sex. In (**B**), genetic correlations among four meiotic recombination phenotypes calculated using individual-level data (MAJA, circles) and summary statistic data (SumHer, triangles), across three countries. Where values are missing, either individual-level data was not accessed (Iceland), or summary statistic estimates of the SNP heritability were not sufficiently different to zero to facilitate a genetic correlation estimate. Error bars represent the 95% credible and confidence intervals, respectively.

**Figure S2:**
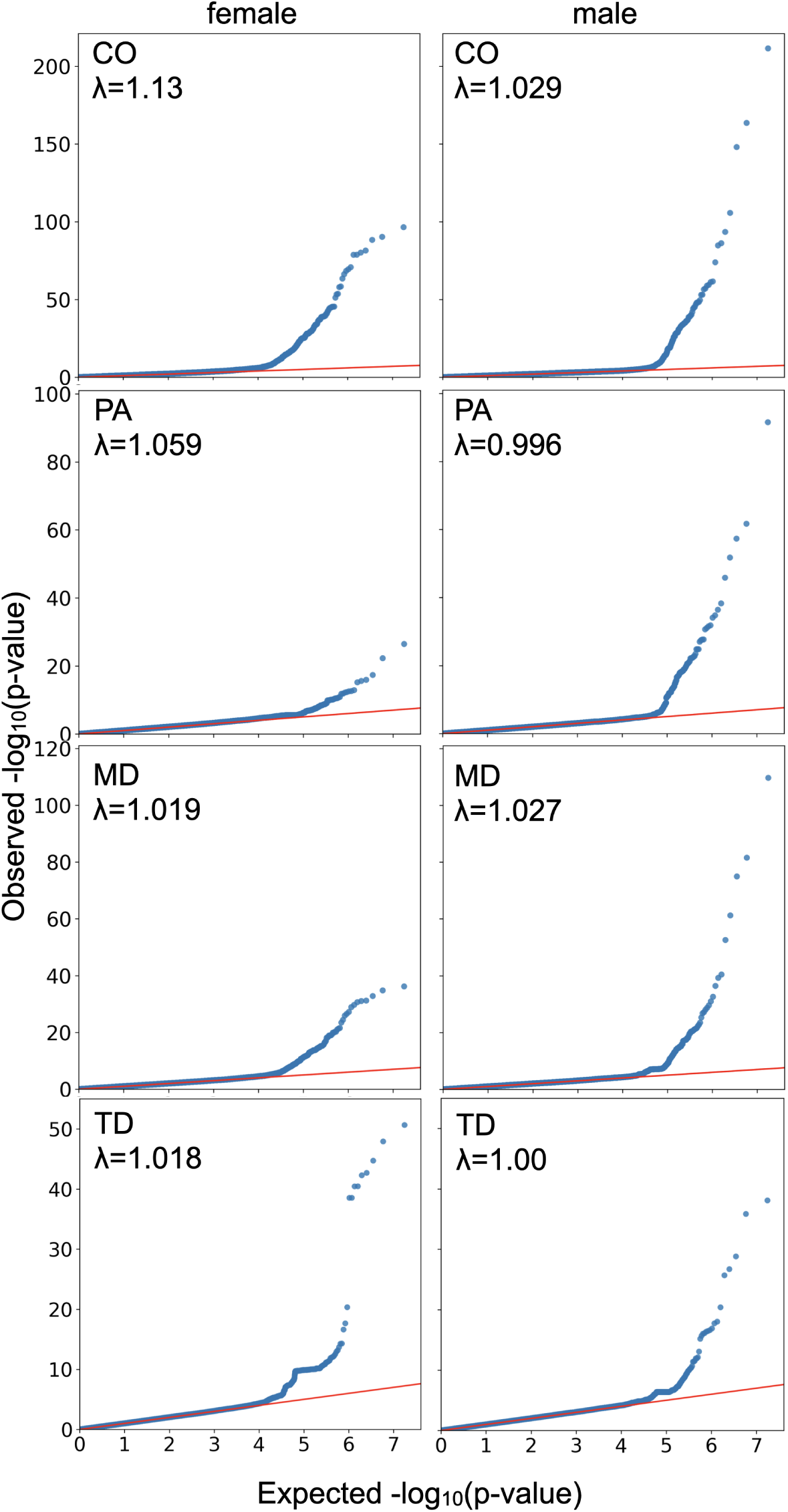
Expected vs observed. 𝜒^2^ **test statistics from the fixed effect meta-analysis.** Values for women (left) and men (right) for recombination cross-over rate (CO), the proportion of single transferred ancestry (PA), mean distance among consecutive cross-over events (MD) and the telomere distance (TD). 𝜆 gives the median divided by the expected median for a 𝜒^2^ distribution with 1 degree of freedom.

**Figure S3:**
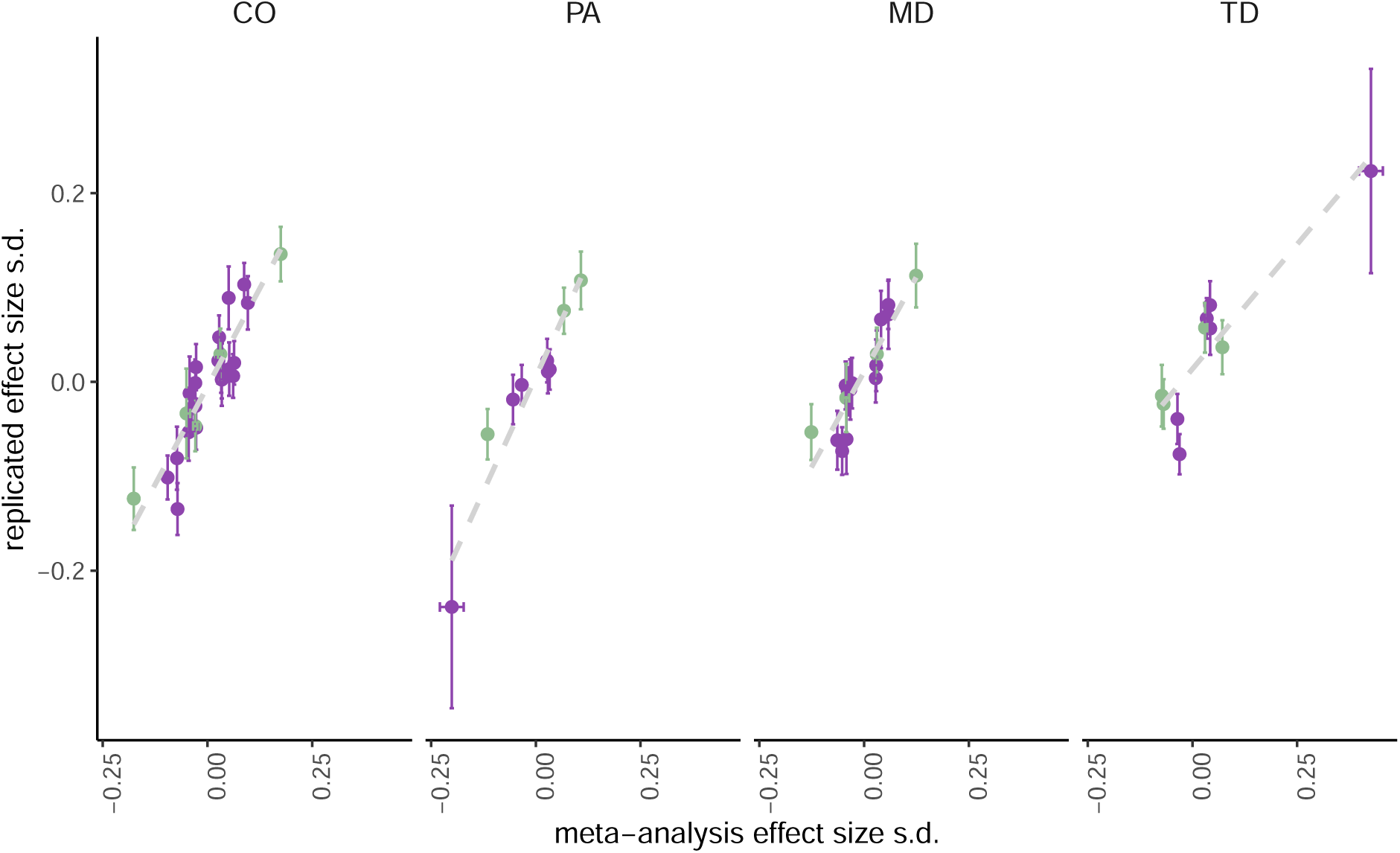
Replication of SNP effect sizes for female and male meiotic recombination in the Generation Scotland cohort plotted against meta-analysis SNP effect sizes for genome-wide significant genetic markers. The panels give the estimates for the measures of recombination cross-over rate (CO), the proportion of single transferred ancestry (PA), the mean distance among consecutive cross-over events (MD) and the telomere distance (TD). Purple and green are female and male meiotic recombination respectively. Error bars give the standard error of the regression coefficient estimates. The slopes of the regression of the replication estimates on the meta-analysis estimates is: 0.830 (0.082 SE, t-statistic=10.09, p-value=2.24 × 10^−14^) for CO; 0.795 (0.122 SE, t-statistic=6.51, p-value< 1.95 × 10^−08^) for MD, 0.967 (0.107 SE, t-statistic=9.01, p-value=1.31 × 10^−12^) for PA; and 0.550 (0.066 SE, t-statistic=8.33, p-value=1.70 × 10^−11^) for TD.

**Figure S4:**
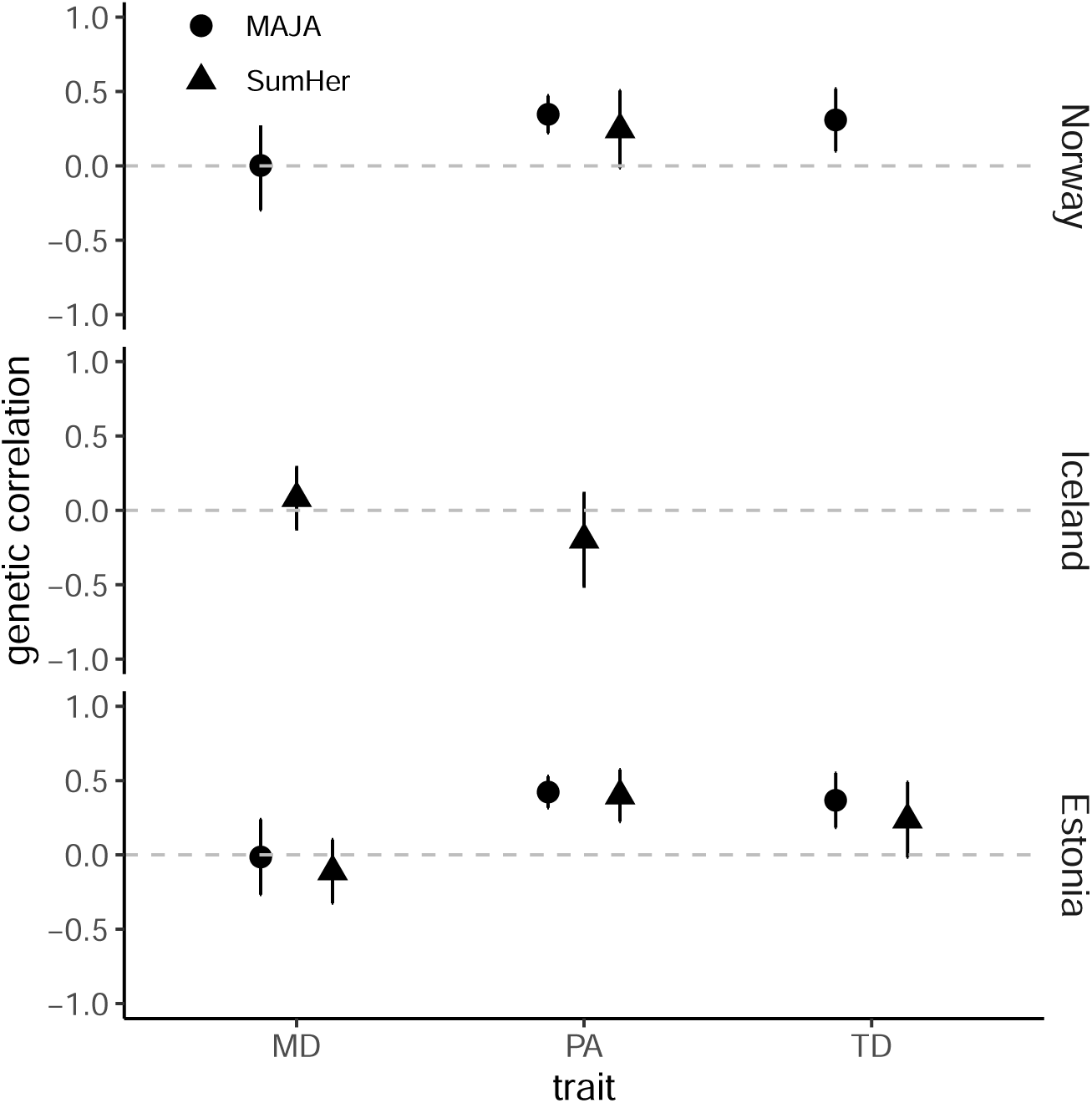
Between-sex genetic correlation for recombination phenotypes PA, MD, and TD across countries. Genetic correlations are calculated using individual-level data (MAJA, circles) and summary statistic data (SumHer, triangles), across Norway, Iceland and Estonia. Where values are missing, either individual-level data was not accessed (Iceland), or summary statistic estimates of the SNP heritability were not sufficiently different to zero to facilitate a genetic correlation estimate. Error bars represent the 95% credible and confidence intervals, respectively.

**Figure S5:**
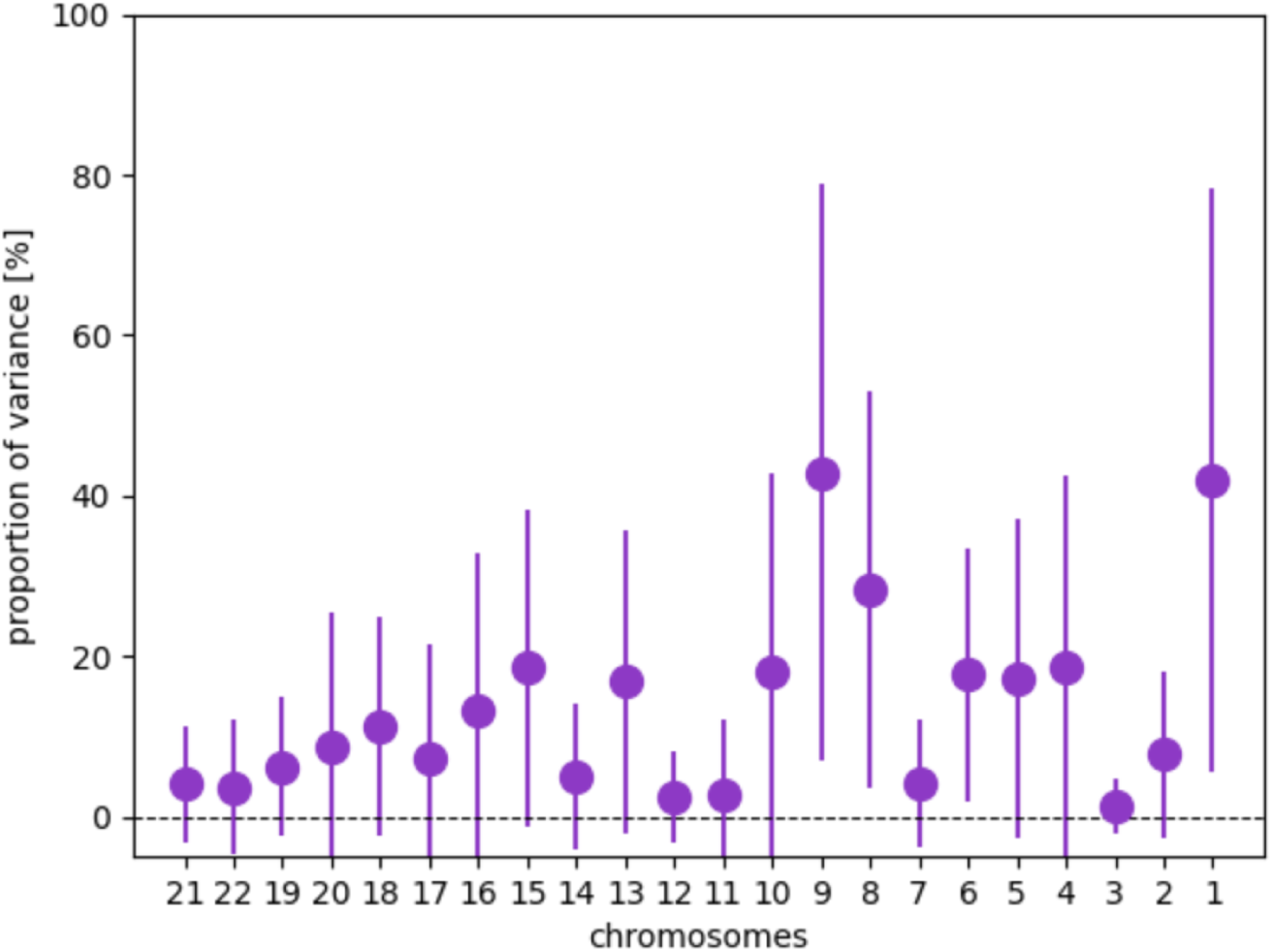
Polygenicity of female recombination rate. Female CO was calculated in the Estonian Biobank (EstBB) 22 times, each time for all chromosomes other than a focal chromosome. We ran 22 models, regressing each CO measure onto two groups of covariates jointly, the genotypes of the focal chromosome and the genotypes of all other chromosomes. For example for chromosome 1, the double strand breaks occurring on chromosomes 2-22 were counted and regressed onto two groups of covariates jointly, the genotypes of chromosome 1 and the genotypes of chromosomes 2-22. Each point in the plot is the proportion of variance attributable to the genotypes of the focal chromosome, with error bars representing the 95% credible intervals. Thus each point is an indication of the degree to which SNPs on a specific chromosome contribute to the variation in CO observed in other chromosomes. The chromosomes shown on the x-axis are ordered by chromosome length.

**Figure S6:**
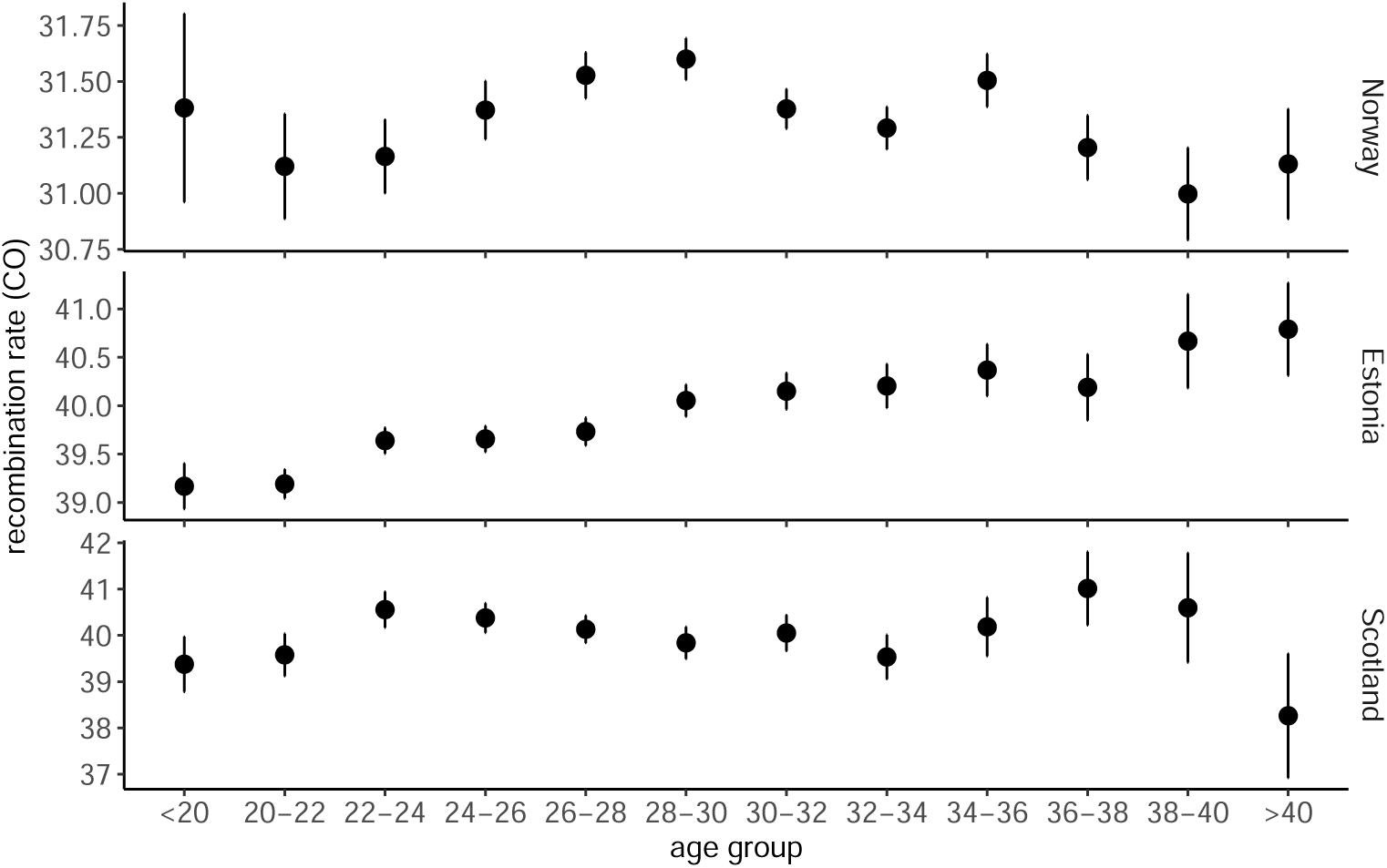
Relationship of female recombination rate and age. Mean female recombination rate (CO) across age groups within each study. Error bars give the SE of the estimate.

**Figure S7:**
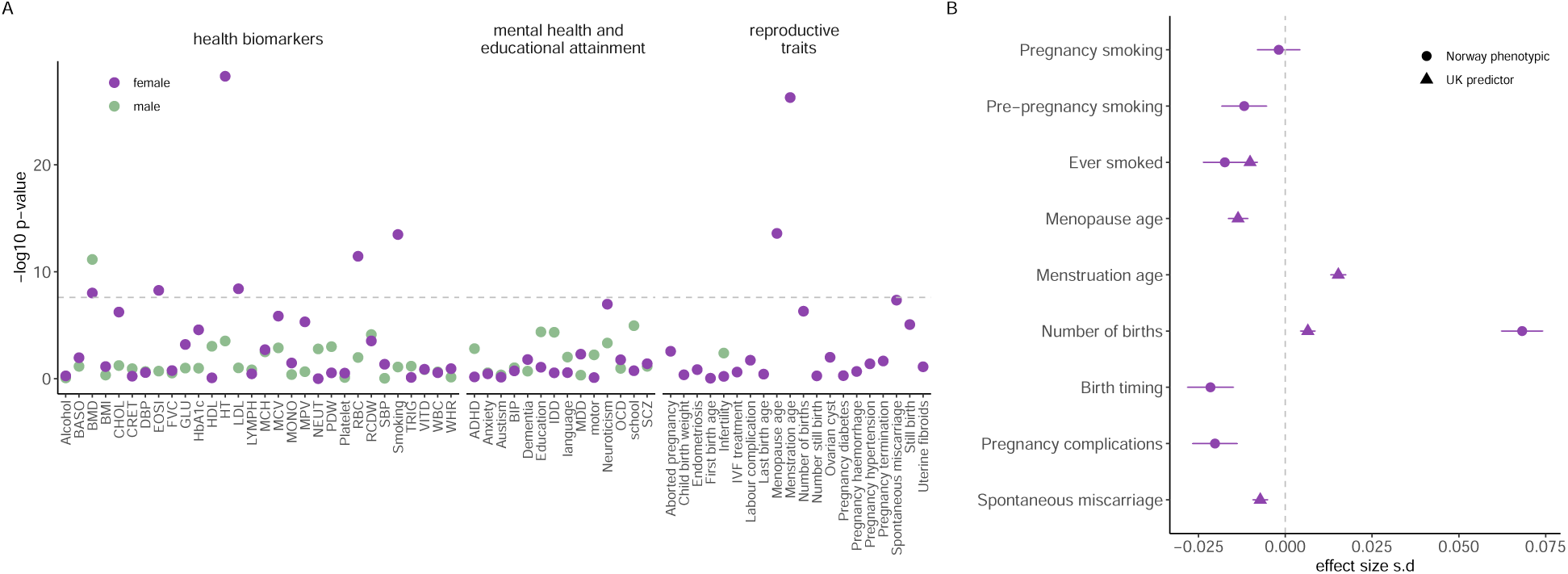
Relationship of meiotic recombination to other phenotypes. In (**A**), each point depicts the −𝑙𝑜𝑔_10_ of the p-value for cross-trait prediction of each trait, where the polygenic risk score for CO in females and males is used to predict that trait, within the UK Biobank study. The dashed gray line represents the multiple testing-corrected p-value threshold of 5 · 10^−6^. In (**B**), for the measure of female recombination cross-over rate (CO), we show the traits for which the CO polygenic risk score is significantly associated in the hold-out UK Biobank sample (triangles) alongside the phenotypic correlation of the most similar traits within the Norway cohort (circles). All estimates are on the correlation scale, as standardized trait values are regressed onto standardized values of the polygenic risk score. Birth timing refers to a binary indicator of whether a pregnancy was conceived more than 12 months from the birth of the last child. Pregnancy complications refers to a binary indicator of self-reported occurrence of complications during pregnancy. Error bars give the 95% intervals of the estimates.

## A Supplementary Tables

**Table S1:**
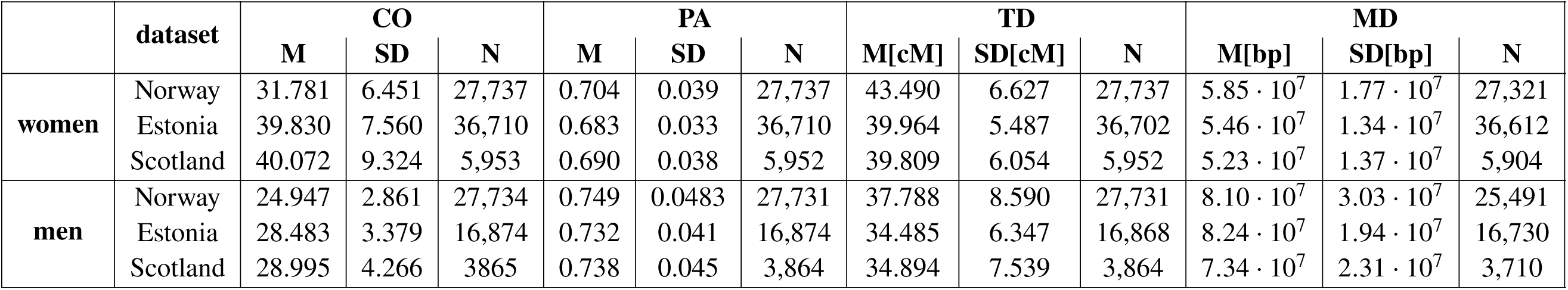
Recombination phenotypes across cohorts for women and men. Phenotypic mean (M), standard deviation (SD) and number of parent-offspring pairs (N) for recombination rate (CO), proportion of ancestry (PA), telomere distance (TD) in cM and mean distance (MD) in bp in Norway, Estonia and Scotland.

**Table S2:** Number of genome-wide significant SNPs for each DNA region across all traits. The association analyses were performed separately in females (f) and males (m) and jointly (j) for both sexes. Four traits were considered: recombination rate (CO), the proportion of single transferred ancestry (PA), the telomere distance (TD) and the mean distance among consecutive cross-over events (MD). Chr, chromosome number; Position, base-pair region. Grey boxes, traits for which a focal SNP and surrounding DNA region was selected from the joint GCTA-cojo analysis.

| Chr | Position | Meta-analysis |  |  |  |  |  |  |  |  |  |  |  |
| --- | --- | --- | --- | --- | --- | --- | --- | --- | --- | --- | --- | --- | --- |
|  |  | CO |  |  | PA |  |  | MD |  |  | TD |  |  |
|  |  | j | f | m | j | f | m | j | f | m | j | f | m |
| 1 | 6701978 : 6784464 | 1 | 3 | 0 |  |  |  |  |  |  |  |  |  |
| 1 | 75841510 : 76259146 | 1 | 4 | 0 |  |  |  | 0 | 1 | 0 |  |  |  |
| 2 | 54416818 : 54508020 | 0 | 5 | 0 |  |  |  | 0 | 0 | 2 | 6 | 3 | 0 |
| 3 | 37053568 | 1 | 1 | 0 |  |  |  |  |  |  |  |  |  |
| 4 | 722812 : 1417603 | 112 | 124 | 155 | 32 | 12 | 96 | 48 | 39 | 110 | 11 | 0 | 34 |
| 4 | 99080544 : 113607978 | 0 | 0 | 1 |  |  |  |  |  |  |  |  |  |
| 5 | 122535307 |  |  |  |  |  |  |  |  |  | 3 | 1 | 0 |
| 5 | 176929582 |  |  |  |  |  |  | 0 | 1 | 0 |  |  |  |
| 6 | 26380032 : 32201279 | 35 | 68 | 0 | 0 | 5 | 0 |  |  |  |  |  |  |
| 10 | 135348026 : 135365335 | 0 | 0 | 1 |  |  |  |  |  |  |  |  |  |
| 12 | 133488642 : 133818020 |  |  |  |  |  |  |  |  |  | 2 | 4 | 0 |
| 14 | 20728125 : 20873524 | 37 | 41 | 0 | 11 | 0 | 0 | 28 | 23 | 0 | 0 | 0 | 1 |
| 14 | 60779884 : 61573059 | 13 | 39 | 9 | 2 | 8 | 0 | 0 | 26 | 3 | 36 | 139 | 0 |
| 14 | 91806588 : 92151515 | 69 | 89 | 0 | 0 | 16 | 0 | 26 | 45 | 0 | 6 | 10 | 0 |
| 16 | 1848748 : 1972548 | 0 | 7 | 0 |  |  |  | 0 | 5 | 0 |  |  |  |
| 16 | 74980490: 75005197 | 0 | 2 | 0 |  |  |  |  |  |  |  |  |  |
| 17 | 37078880 : 44888742 | 50 | 126 | 0 | 0 | 1 | 0 | 6 | 53 | 0 | 228 | 28 | 0 |
| 18 | 12928541 : 13164588 | 1 | 6 | 0 |  |  |  |  |  |  |  |  |  |
| 18 | 74626997 : 74691452 | 0 | 4 | 0 |  |  |  | 0 | 3 | 0 |  |  |  |
| 19 | 10149516 : 13220749 | 4 | 14 | 0 | 0 | 9 | 0 |  |  |  | 11 | 19 | 0 |
| 19 | 20258708 : 22631153 | 2 | 3 | 0 |  |  |  | 3 | 3 | 0 |  |  |  |
| 20 | 1175527 : 7366771 | 3 | 3 | 1 | 12 | 0 | 13 | 1 | 1 | 0 | 4 | 0 | 13 |
| 20 | 56096508: 61327966 | 0 | 1 | 0 |  |  |  | 0 | 0 | 1 |  |  |  |
| 22 | 45828546 : 50594835 | 0 | 2 | 0 |  |  |  | 6 | 2 | 0 |  |  |  |

**Table S3:** Number of genome-wide significant SNPs for each DNA region across all traits jointly fit with the ’cojo’ algorithm of the GCTA software. The association analyses were performed separately in females (f) and males (m) and jointly (j) for both sexes. Four traits were considered: recombination rate (CO), the proportion of single transferred ancestry (PA), the telomere distance (TD) and the mean distance among consecutive cross-over events (MD). Chr, chromosome number; Position, base-pair region. Grey boxes, traits and biological sexes for which a focal SNP and surrounding DNA region was selected from the joint GCTA-cojo analysis.

| Chr | Position | GCTA - cojo SNPs |  |  |  |  |  |  |  |  |  |  |  |
| --- | --- | --- | --- | --- | --- | --- | --- | --- | --- | --- | --- | --- | --- |
|  |  | CO |  |  | PA |  |  | MD |  |  | TD |  |  |
|  |  | j | f | m | j | f | m | j | f | m | j | f | m |
| 1 | 6701978 : 6784464 | 2 | 2 |  |  |  |  |  |  |  |  |  |  |
| 1 | 75841510 : 76259146 | 1 | 1 |  |  |  |  |  | 1 |  |  |  |  |
| 2 | 54416818 : 54508020 |  | 9 |  |  |  |  |  |  | 3 | 3 | 3 |  |
| 3 | 37053568 | 2 | 3 |  |  |  |  |  |  |  |  |  |  |
| 4 | 722812 : 1417603 | 17 | 16 | 20 | 15 | 21 | 18 | 16 | 6 | 20 | 5 |  | 16 |
| 4 | 99080544 : 113607978 |  |  | 1 |  |  |  |  |  | 1 |  |  |  |
| 5 | 122535307 |  |  |  |  |  |  |  |  |  | 1 | 1 |  |
| 5 | 176929582 |  |  |  |  |  |  |  | 1 |  |  |  |  |
| 6 | 26380032 : 32201279 | 9 | 17 |  |  | 11 |  |  |  |  |  |  |  |
| 10 | 135348026 : 135365335 |  |  | 2 |  |  |  |  |  |  |  |  |  |
| 12 | 133488642 : 133818020 |  |  |  |  |  |  |  |  |  | 7 | 12 |  |
| 14 | 20728125 : 20873524 | 5 | 10 |  | 4 |  |  | 9 | 8 |  |  |  | 2 |
| 14 | 60779884 : 61573059 | 2 | 2 |  |  |  |  |  |  |  |  |  |  |
| 14 | 91806588 : 92151515 | 10 | 9 |  |  | 1 |  | 12 | 13 |  | 4 | 6 |  |
| 16 | 1848748 : 1972548 |  | 2 |  |  |  |  |  | 6 |  |  |  |  |
| 16 | 74980490 : 75005197 |  | 1 |  |  |  |  |  |  |  |  |  |  |
| 17 | 37078880 : 44888742 | 26 | 25 |  |  | 1 |  | 4 | 20 |  | 11 | 7 |  |
| 18 | 12928541 : 13164588 | 4 | 5 |  |  |  |  |  |  |  |  |  |  |
| 18 | 74626997 : 74691452 |  | 5 |  |  |  |  |  | 8 |  |  |  |  |
| 19 | 10149516 : 13220749 | 7 | 11 |  |  | 1 |  |  |  |  | 2 | 5 |  |
| 19 | 20258708 : 22631153 | 5 | 1 |  |  |  |  |  |  |  |  |  |  |
| 20 | 1175527 : 7366771 | 8 | 2 |  | 10 |  | 11 | 1 | 1 |  | 1 |  | 5 |
| 20 | 56096508 : 61327966 |  | 1 |  |  |  |  |  |  | 3 |  |  |  |
| 22 | 45828546 : 50594835 |  | 5 |  |  |  |  | 6 | 5 |  |  |  |  |

**Table S4:** Fine-mapped SNPs for each DNA region across all traits using the fine-mapping algorithm of the SuSieX software. The association analyses were performed separately in females (f) and males (m). Four traits were considered: recombination rate (CO), the proportion of single transferred ancestry (PA), the telomere distance (TD) and the mean distance among consecutive cross-over events (MD). Chr, chromosome number; Position, base-pair region. ’rs’ identifiers are focal SNPs of a credible set with posterior inclusion probability ≥ 0.95 and n gives the number of SNPs within each credible set.

| Chr | Position | SuSieX - fine-mapped SNPs |  |  |  |  |  |  |  |  |  |  |  |
| --- | --- | --- | --- | --- | --- | --- | --- | --- | --- | --- | --- | --- | --- |
|  |  | CO |  |  | PA |  |  | MD |  |  | TD |  |  |
|  |  | n | f | m | n | f | m | n | f | m | n | f | m |
| 1 | 6701978 : 6784464 | 1<br>3 | rs112793972<br>rs72865913 | rs112793972<br>rs72865913 |  |  |  |  |  |  |  |  |  |
| 1 | 75841510 : 76259146 | 1 | rs6686883 | - |  |  |  | 1<br>1 | rs6686883<br>rs5745459 | -<br>- |  |  |  |
| 2 | 54416818 : 54508020 | 1 | rs10205866 | rs10205866 |  |  |  | 1<br>1<br>1 | rs6740641<br>rs62139199<br>rs12618994 | rs6740641<br>rs62139199<br>rs12618994 | 2 | rs10205866 | rs10205866 |
| 3 | 37053568 | - | - | - |  |  |  |  |  |  |  |  |  |
| 4 | 722812 : 1417603 | 2 | rs3816474 | rs3816474 | 1 | rs2290409 | rs2290409 | 1 | rs2290409 | rs2290409 | 1 | - | rs2290409 |
| 4 | 99080544 : 113607978 | 3<br>1<br>5 | rs973126<br>rs2522434<br>rs6844767 | rs973126<br>rs2522434<br>rs6844767 |  |  |  |  |  |  |  |  |  |
| 5 | 122535307 |  |  |  |  |  |  |  |  |  | 1 | rs31319 | rs31319 |
| 5 | 122535307 |  |  |  |  |  |  |  |  |  | 1 | rs31318 | rs31318 |
| 5 | 176929582 |  |  |  |  |  |  | 9<br>1 | rs60780889<br>rs6883391 | rs60780889<br>rs6883391 |  |  |  |
| 6 | 26380032 : 32201279 | - | - | - | - | - | - |  |  |  |  |  |  |
| 10 | 135348026 : 135365335 | 1<br>1 | rs11017846<br>rs1055043 | rs11017846<br>rs1055043 |  |  |  |  |  |  |  |  |  |
| 12 | 133488642 : 133818020 |  |  |  |  |  |  |  |  |  | - | - | - |
| 14 | 20728125 : 20873524 | 1 | rs1953549 | rs1953549 |  |  |  | 2 | rs1953549 | rs1953549 | 1<br>2<br>1<br>1 | rs2781366<br>rs4981975<br>rs1713419<br>rs1713418 | rs2781366<br>rs4981975<br>rs1713419<br>rs1713418 |
| 14 | 60779884 : 61573059 | 1 | rs1254251 | rs1254251 | 1 | rs1254251 | - | 1 | rs1254251 | rs1254251 | 2<br>1 | rs1254252<br>rs1254251 | rs1254252<br>rs1254251 |
| 14 | 91806588 : 92151515 | 2 | rs1961540 | - | 1<br>1 | rs35144173<br>rs34255419 | rs35144173<br>rs34255419 | 2 | rs1961540 | - | 2 | rs1961540 | - |
| 16 | 1848748 : 1972548 | 1 | rs1623559 | rs1623559 |  |  |  | 5 | rs1623559 | - |  |  |  |
| 16 | 74980490 : 75005197 | 1<br>1 | rs143068306<br>rs143235261 | rs143068306<br>rs143235261 |  |  |  |  |  |  |  |  |  |
| 17 | 37078880 : 44888742 | 3 | rs3785884 | - | 1<br>1 | rs8071912<br>rs1598765 | rs8071912<br>rs1598765 | - | - | - | 3 | rs3785884 | - |
| 18 | 12928541 : 13164588 | 1<br>1 | rs1474043<br>rs4995995 | -<br>- |  |  |  |  |  |  |  |  |  |
| 18 | 74626997 : 74691452 | 1<br>1 | rs4640270<br>rs8084848 | rs4640270<br>rs8084848 |  |  |  | 1<br>1 | rs4640270<br>rs8084848 | rs4640270<br>rs8084848 |  |  |  |

| Table S6 continued |  |  |  |  |  |  |  |  |  |  |  |  |  |
| --- | --- | --- | --- | --- | --- | --- | --- | --- | --- | --- | --- | --- | --- |
| 19 | 10149516 : 13220749 | 1<br>1<br>1 | rs34275421<br>rs28642201<br>rs8105074 | -<br>rs28642201<br>rs8105074 | 2<br>1<br>1<br>1<br>1<br>1<br>13<br>1 | rs184545813<br>rs139062350<br>rs117680228<br>rs189601197<br>rs117547594<br>rs116857070<br>rs11673414<br>rs7254518 | rs184545813<br>rs139062350<br>rs117680228<br>rs189601197<br>rs117547594<br>rs116857070<br>rs11673414<br>rs7254518 |  |  |  | 1<br>1<br>2<br>1 | rs117680228<br>rs139062350<br>rs184545813<br>rs189601197 | -<br>rs139062350<br>rs184545813<br>rs189601197 |
| 19 | 20258708 : 22631153 | 1<br>2<br>1<br>1 | rs1230295<br>rs12977311<br>rs12609595<br>rs1230305 | rs1230295<br>rs12977311<br>rs12609595<br>rs1230305 |  |  |  | 1<br>1<br>1<br>1 | rs74180875<br>rs34210970<br>rs12977553<br>- | rs74180875<br>rs34210970<br>rs12977553<br>rs74180876 |  |  |  |
| 20 | 1175527 : 7366771 | 1<br>1 | rs450739<br>rs203535 | rs450739<br>rs203535 | 1 | - | rs450739 | 1<br>1 | rs450739<br>rs203535 | rs450739<br>rs203535 | 1 | - | rs450739 |
| 20 | 56096508 : 61327966 | 4<br>1 | rs79080761<br>rs1040782 | rs79080761<br>rs1040782 |  |  |  | - | - | - |  |  |  |
| 22 | 45828546 : 50594835 | - | - | - |  |  |  | 1<br>1<br>1 | rs11702941<br>rs78947992<br>rs11703201 | rs11702941<br>rs78947992<br>rs11703201 |  |  |  |

**Table S5:**
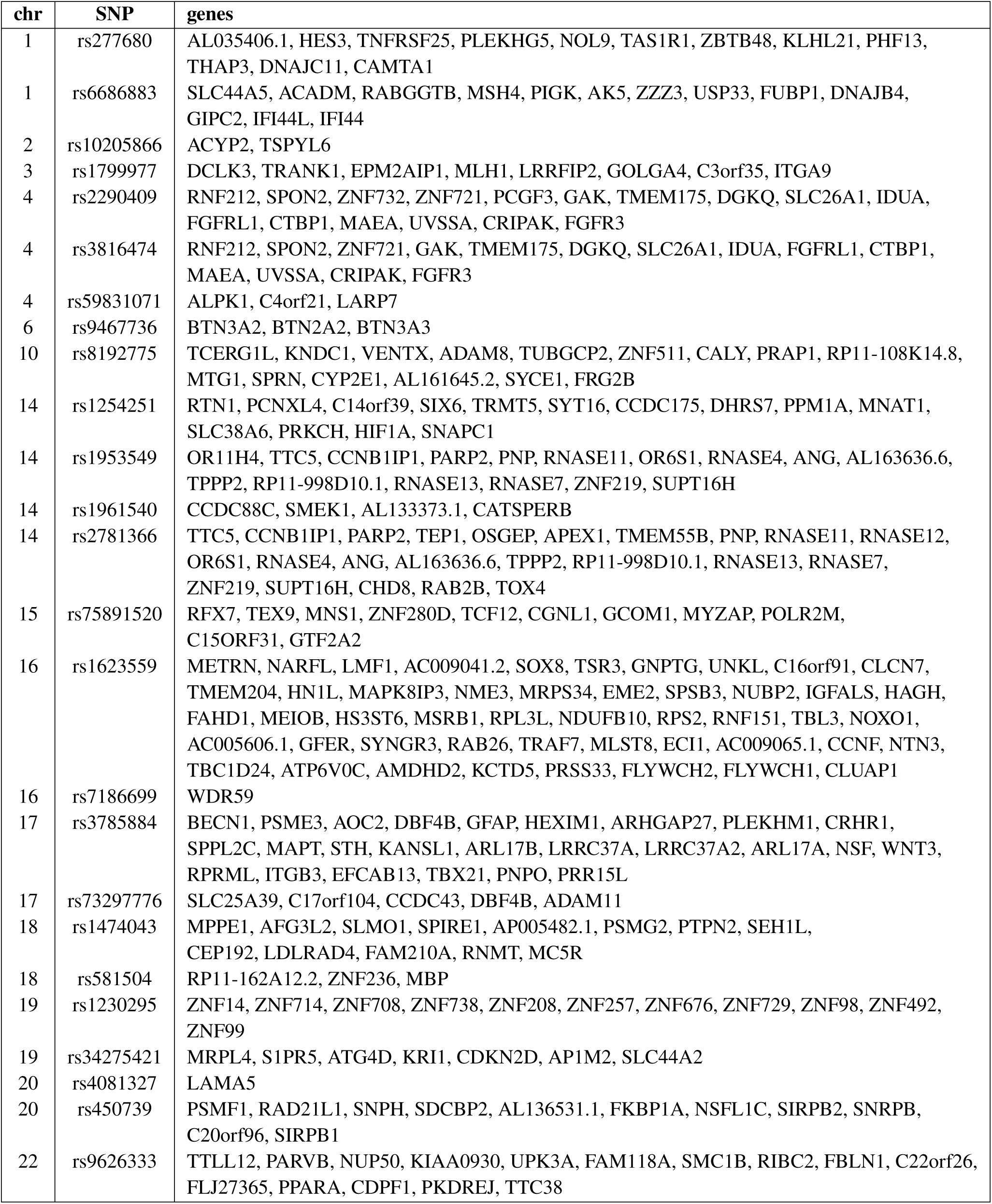
Genes mapped to significant SNPs found for CO. Chromosome number (chr), rsid identifier (SNP), and genes identified with FUMA for leading associated SNPs within each region for CO given in Table 1.

**Table S6:** Genes mapped to significant SNPs found for PA. Chromosome number (chr), rsid identifier (SNP), and genes identified with FUMA for leading associated SNPs within each region for PA given in Table 1.

| chr | SNP | genes |
| --- | --- | --- |
| 4 | rs2290409 | ZNF732, ZNF721, PCGF3, GAK, TMEM175, DGKQ, SLC26A1, IDUA, FGFR1, RNF212, SPON2, CTBP1, MAEA, UVSSA, CRIPAK, NKX1-1, FGFR3 |
| 4 | rs3816474 | ZNF721, GAK, TMEM175, DGKQ, SLC26A1, IDUA, FGFR1, RNF212, SPON2, CTBP1, MAEA, UVSSA, CRIPAK, FGFR3, ZNF732, ATP5I, MYL5, NKX1-1 |
| 14 | rs1254251 | CCDC175, PCNXL4, DHRS7, PPM1A, C14orf39, SIX6, MNAT1, TRMT5, SLC38A6, HIF1A, SNAPC1 |
| 14 | rs1285841 | GPR68, CCDC88C, SMEK1, AL133373.1, CATSPERB, TC2N, FBLN5, TRIP11, NDUFB1, CPSF2, RIN3 |
| 19 | rs4987202 | ZNF878, GCDH, SYCE2, CALR, RAD23A, GADD45GIP1, STX10, IER2 |
| 20 | rs450739 | C20orf96, PSMF1, RAD21L1, SNPH, FKBP1A, SIRPB1 |

**Table S7:** Genes mapped to significant SNPs found for TD. Chromosome number (chr), rsid identifier (SNP), and genes identified with FUMA for leading associated SNPs within each region for TD given in Table 1.

| chr | SNP | genes |
| --- | --- | --- |
| 2 | rs10205866 | GPR75-ASB3, ACYP2, TSPYL6, C2orf73, SPTBN1, EML6, CLHC1, RPS27A, |
| 4 | rs2290409 | ZNF732, PCGF3, GAK, TMEM175, DGKQ, SLC26A1, IDUA, FGFR1, RNF212, CCDC104, SPON2, CTBP1, MAEA, UVSSA, CRIPAK |
| 4 | rs3816474 | GAK, TMEM175, DGKQ, SLC26A1, IDUA, FGFR1, RNF212, SPON2, CTBP1, MAEA, UVSSA, CRIPAK |
| 5 | rs143073807 | LOX, SNCAIP, SNX2, SNX24, PPIC, PRDM6, CEP120, CSNK1G3 |
| 12 | rs36179555 | GOLGA3, ZNF605, ZNF84, ZNF140, ZNF891, ZNF10, ZNF268, CTD-2140B24.4, AC226150.4, ANHX |
| 14 | rs1961540 | NRDE2, GPR68, CCDC88C, SMEK1, AL133373.1, CATSPERB, TC2N, TRIP11, ATXN3, NDUFB1, CPSF2, RIN3 |
| 14 | rs2781366 | TTC5, CCNB1IP1, PARP2, TEP1, OSGEP, APEX1, TMEM55B, PNP, RNASE11, RNASE12, OR6S1, RNASE4, ANG, AL163636.6, RNASE1, RNASE2, RNASE7, ZNF219, SUPT16H |
| 14 | rs8015263 | CCDC175, RTN1, PCNXL4, DHRS7, PPM1A, C14orf39, SIX6, SIX1, MNAT1, TRMT5, PRKCH, TMEM30B, HIF1A, SNAPC1, SYT16, KCNH5 |
| 17 | rs3785884 | GPS2, GID4, RAB34, MPP3, FZD2, CCDC43, DBF4B, GJC1, HIGD1B, EFTUD2, CCDC103, FAM187A, GFAP, DCAKD, NMT1, PLCD3, ACBD4, HEXIM1, HEXIM2, SPATA32, ARHGAP27, PLEKHM1, CRHR1, SPPL2C, MAPT, STH, LRRC37A, LRRC37A2, ARL17A, NSF, WNT3, WNT9B, GOSR2, RP11-156P1.2, MYL4, ITGB3, EFCAB13, KPNB1, TBKBP1, TBX21, SP6, HOXB1, HOXB2, HOXB7, MKS1, LRRC37A3, C17orf58, LGALS3BP, ARL17B, CDC27, KANSL1, FMNL1, RPRML |
| 19 | rs4987202 | ZNF878, GCDH, SYCE2, CALR, RAD23A, GADD45GIP1 |
| 20 | rs450739 | C20orf96, PSMF1, RAD21L1, SNPH, FKBP1A, SIRPB1 |

**Table S8:** Genes mapped to significant SNPs found for MD. Chromosome number (chr), rsid identifier (SNP), and genes identified with FUMA for leading associated SNPs within each region for MD given in Table 1.

| chr | SNP | genes |
| --- | --- | --- |
| 1 | rs6686883 | CRYZ, LHX8, SLC44A5, ACADM, RABGGTB, MSH4, ST6GALNAC3, ST6GALNAC5, PIGK, AK5, USP33, FUBP1, DNAJB4, GIPC2 |
| 2 | rs10205866 | GPR75-ASB3, ACYP2, TSPYL6, C2orf73, SPTBN1, EML6, CLHC1, RPS27A, MTIF2, CCDC104 |
| 4 | rs2290409 | ZNF732, ZNF721, PCGF3, GAK, TMEM175, DGKQ, SLC26A1, IDUA, FGFR1, RNF212, SPON2, CTBP1, MAEA, UVSSA, CRIPAK, NKX1-1, FGFR3 |
| 4 | rs3816474 | ZNF721, GAK, TMEM175, DGKQ, SLC26A1, IDUA, FGFR1, RNF212, SPON2, CTBP1, MAEA, UVSSA, CRIPAK, FGFR3 |
| 5 | rs28712799 | LMAN2, RGS14, F12, GRK6, PRR7, DBN1, PDLIM7, DOK3, DDX41, PHYKPL |
| 14 | rs1254251 | CCDC175, PCNXL4, DHRS7, PPM1A, C14orf39, SIX6, MNAT1, TRMT5, SLC38A6, HIF1A, SNAPC1 |
| 14 | rs1953549 | TTC5, CCNB1IP1, PARP2, TMEM55B, PNP, RNASE11, OR6S1, RNASE4, ANG, AL163636.6, RNASE1, RNASE7, ZNF219, SUPT16H |
| 14 | rs1961540 | CCDC88C, SMEK1, AL133373.1, CATSPERB, TC2N |
| 14 | rs2781366 | TTC5, CCNB1IP1, PARP2, TEP1, RNASE1, RNASE2 |
| 16 | rs1623559 | METRNL, NARFL, LMF1, AC009041.2, SOX8, TPSG1, TPSD1, TSR3, GNPTG, UNKL, C16orf91, CLCN7, IFT140, TMEM204, CRAMP1L, HN1L, MAPK8IP3, NME3, MRPS34, EME2, SPSB3, NUBP2, IGFALS, HAGH, FAHD1, MEIOB, HS3ST6, MSRB1, RPL3L, NDUFB10, RPS2, RNF151, TBL3, NOXO1, GFER, SYNGR3, ZNF598, PKD1, RAB26, TRAF7, MLST8, E4F1, ECI1, AC009065.1, CCNF, NTN3, TBC1D24, ATP6V0C, AMDHD2, KCTD5, PRSS27, AC005606.1, PRSS33, FLYWCH2, FLYWCH1, CLUAP1 |
| 17 | rs73297776 | SLC25A39, FZD2, C17orf104, CCDC43, DBF4B, ADAM11, GJC1, HIGD1B, EFTUD2, DCAKD, PLCD3, ARHGAP27, LRRC37A2, LRRC37A3 |
| 18 | rs4640270 | FBXO15, TIMM21, C18orf63, FAM69C, ZNF407, LINC00908, RP11-162A12.2, ZNF236, MBP, GALR1, SALL3 |
| 20 | rs450739 | C20orf96, PSMF1, RAD21L1, SNPH, FKBP1A, NSFL1C, SIRPB2, SIRPB1, SNRPB |
| 20 | rs6025604 | RTFDC1, RAE1, MTRNR2L3, CTCFL, PCK1, ZBP1, C20orf85, ANKRD60, PPP4R1L, RAB22A, VAPB, MGC4294, STX16, STX16-NPEPL1, NPEPL1, GNAS, CTSZ, TUBB1, ATP5E, SLMO2, ZNF831, EDN3, PHACTR3 |
| 22 | rs11913200 | PARVB, ARHGAP8, NUP50, KIAA0930, UPK3A, FAM118A, SMC1B, RIBC2, FBLN1, FLJ27365, PPARA, CDPF1, PKDREJ, TTC38 |

**Table S9:**
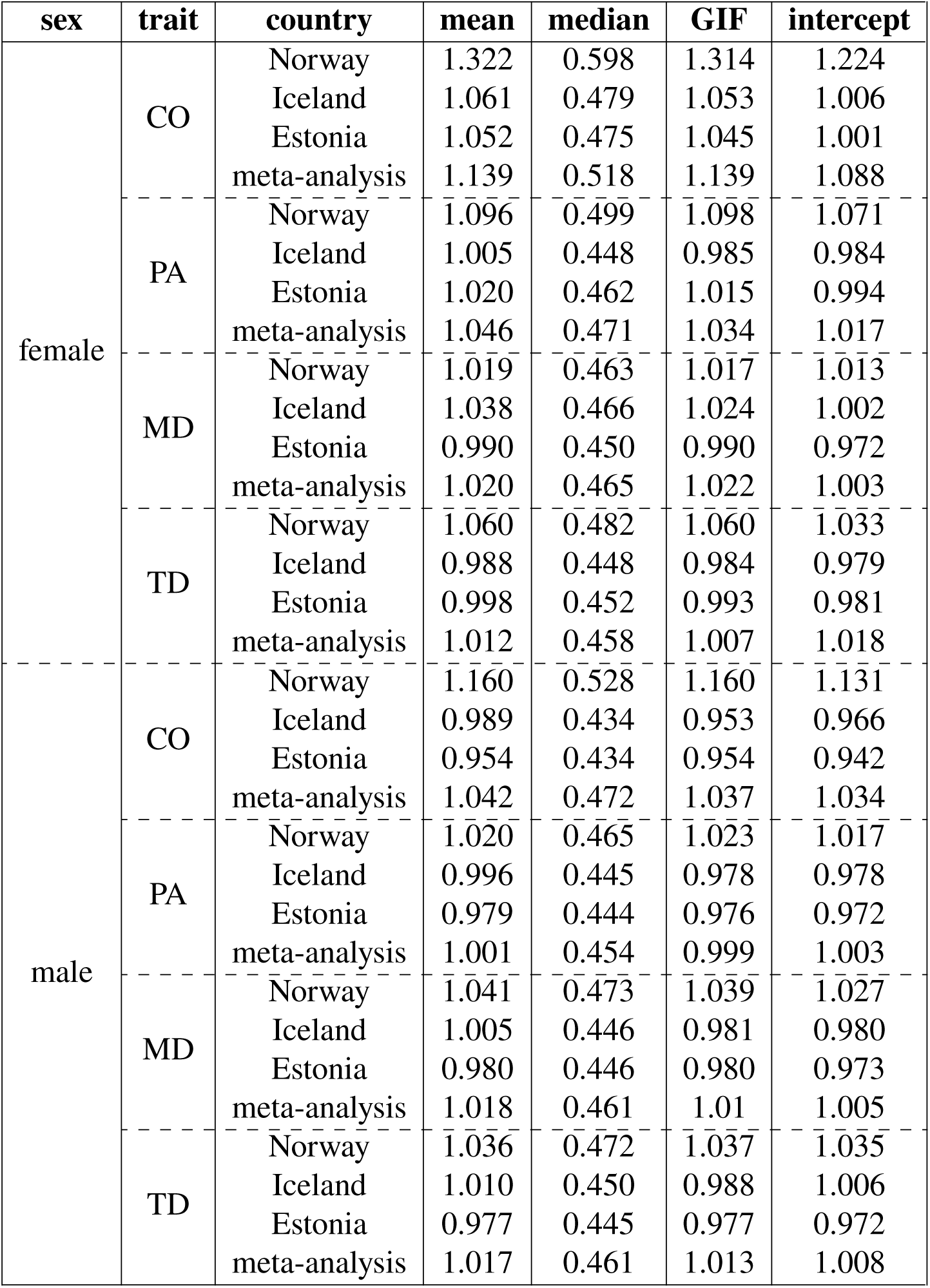
Mean, median, genomic inflation factor (GIF) and SumHer intercept for the test statistic. 𝜒^2^ **values across SNPs from the mixed-linear model association analysis of each cohort.** GIF refers to genomic inflation factor, the median 𝜒^2^ value divided by the expectation. Intercept refers to the SumHer intercept estimate. Four meiotic recombination phenotypes: recombination rate (CO), proportion of ancestry (PA), telomere distance (TD), and mean distance (MD) are analyzed for females and males separately. ”meta-analysis” refers to the fixed-effect inverse variance weighted meta-analysis.

| sex | trait | country | mean | median | GIF | intercept |
| --- | --- | --- | --- | --- | --- | --- |
| female | CO | Norway | 1.322 | 0.598 | 1.314 | 1.224 |
|  |  | Iceland | 1.061 | 0.479 | 1.053 | 1.006 |
|  |  | Estonia | 1.052 | 0.475 | 1.045 | 1.001 |
|  |  | meta-analysis | 1.139 | 0.518 | 1.139 | 1.088 |
|  | PA | Norway | 1.096 | 0.499 | 1.098 | 1.071 |
|  |  | Iceland | 1.005 | 0.448 | 0.985 | 0.984 |
|  |  | Estonia | 1.020 | 0.462 | 1.015 | 0.994 |
|  |  | meta-analysis | 1.046 | 0.471 | 1.034 | 1.017 |
|  | MD | Norway | 1.019 | 0.463 | 1.017 | 1.013 |
|  |  | Iceland | 1.038 | 0.466 | 1.024 | 1.002 |
|  |  | Estonia | 0.990 | 0.450 | 0.990 | 0.972 |
|  |  | meta-analysis | 1.020 | 0.465 | 1.022 | 1.003 |
|  | TD | Norway | 1.060 | 0.482 | 1.060 | 1.033 |
|  |  | Iceland | 0.988 | 0.448 | 0.984 | 0.979 |
|  |  | Estonia | 0.998 | 0.452 | 0.993 | 0.981 |
|  |  | meta-analysis | 1.012 | 0.458 | 1.007 | 1.018 |
| male | CO | Norway | 1.160 | 0.528 | 1.160 | 1.131 |
|  |  | Iceland | 0.989 | 0.434 | 0.953 | 0.966 |
|  |  | Estonia | 0.954 | 0.434 | 0.954 | 0.942 |
|  |  | meta-analysis | 1.042 | 0.472 | 1.037 | 1.034 |
|  | PA | Norway | 1.020 | 0.465 | 1.023 | 1.017 |
|  |  | Iceland | 0.996 | 0.445 | 0.978 | 0.978 |
|  |  | Estonia | 0.979 | 0.444 | 0.976 | 0.972 |
|  |  | meta-analysis | 1.001 | 0.454 | 0.999 | 1.003 |
|  | MD | Norway | 1.041 | 0.473 | 1.039 | 1.027 |
|  |  | Iceland | 1.005 | 0.446 | 0.981 | 0.980 |
|  |  | Estonia | 0.980 | 0.446 | 0.980 | 0.973 |
|  |  | meta-analysis | 1.018 | 0.461 | 1.01 | 1.005 |
|  | TD | Norway | 1.036 | 0.472 | 1.037 | 1.035 |
|  |  | Iceland | 1.010 | 0.450 | 0.988 | 1.006 |
|  |  | Estonia | 0.977 | 0.445 | 0.977 | 0.972 |
|  |  | meta-analysis | 1.017 | 0.461 | 1.013 | 1.008 |

**Table S10:**
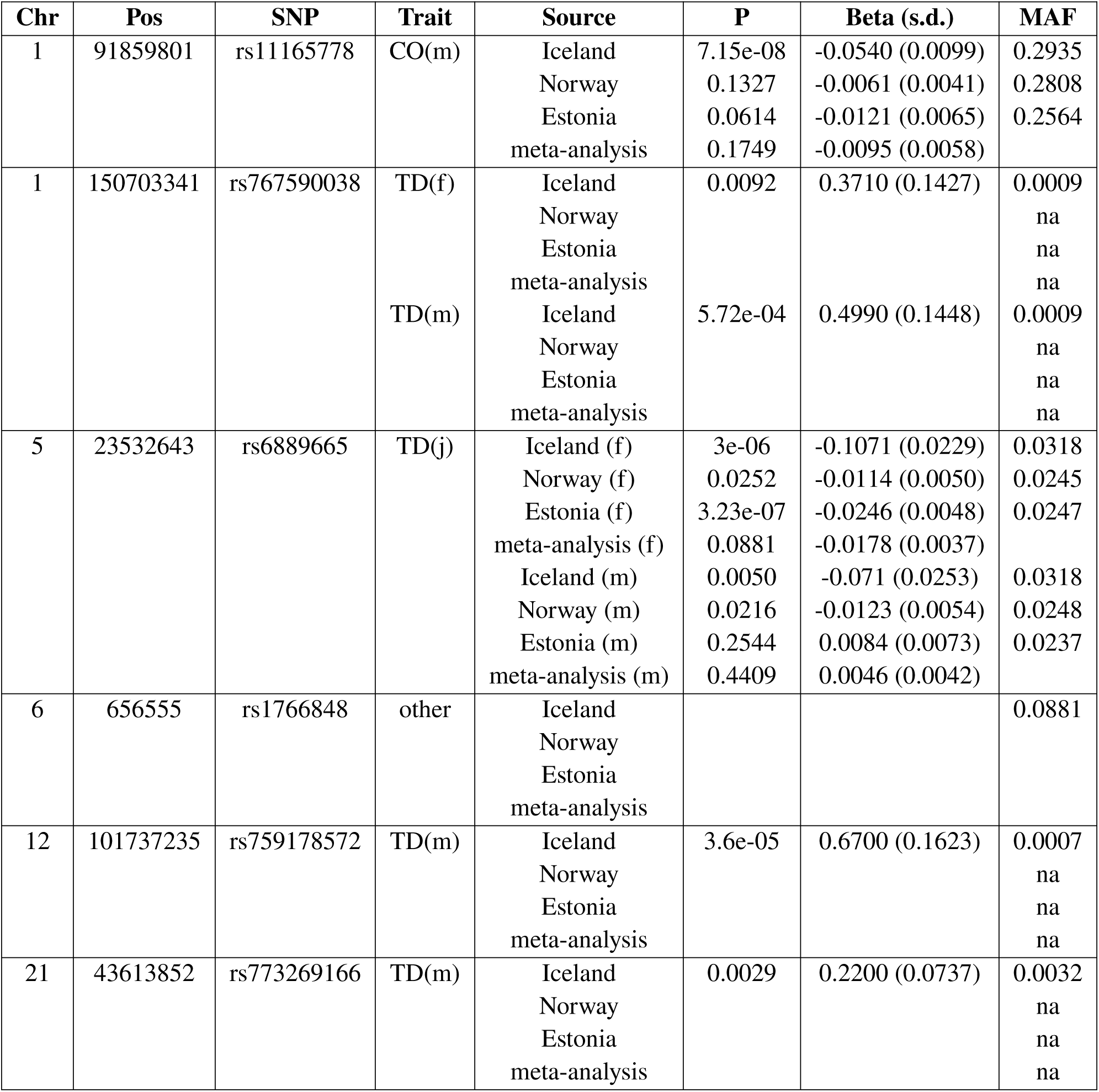
Associated SNPs found in Ref. (***10***)**, not replicated in our meta-analysis.** For each SNP identified by chromosome (Chr) and base pair window (Pos), the p-value (P), the effect sizes (Beta) and minor allele frequency (MAF) are given for the trait (sex) associated within Ref. (*10*) for the different cohorts (Source). If the trait is given as ’other’, the SNP is associated due to other phenotypes that we did not analyze in our study. ’m’ refers to men, ’f’ to female and ’j’ to joint. ’na’ indicates that the SNP was not present after QC in the cohort individuals.

**Table S11:**
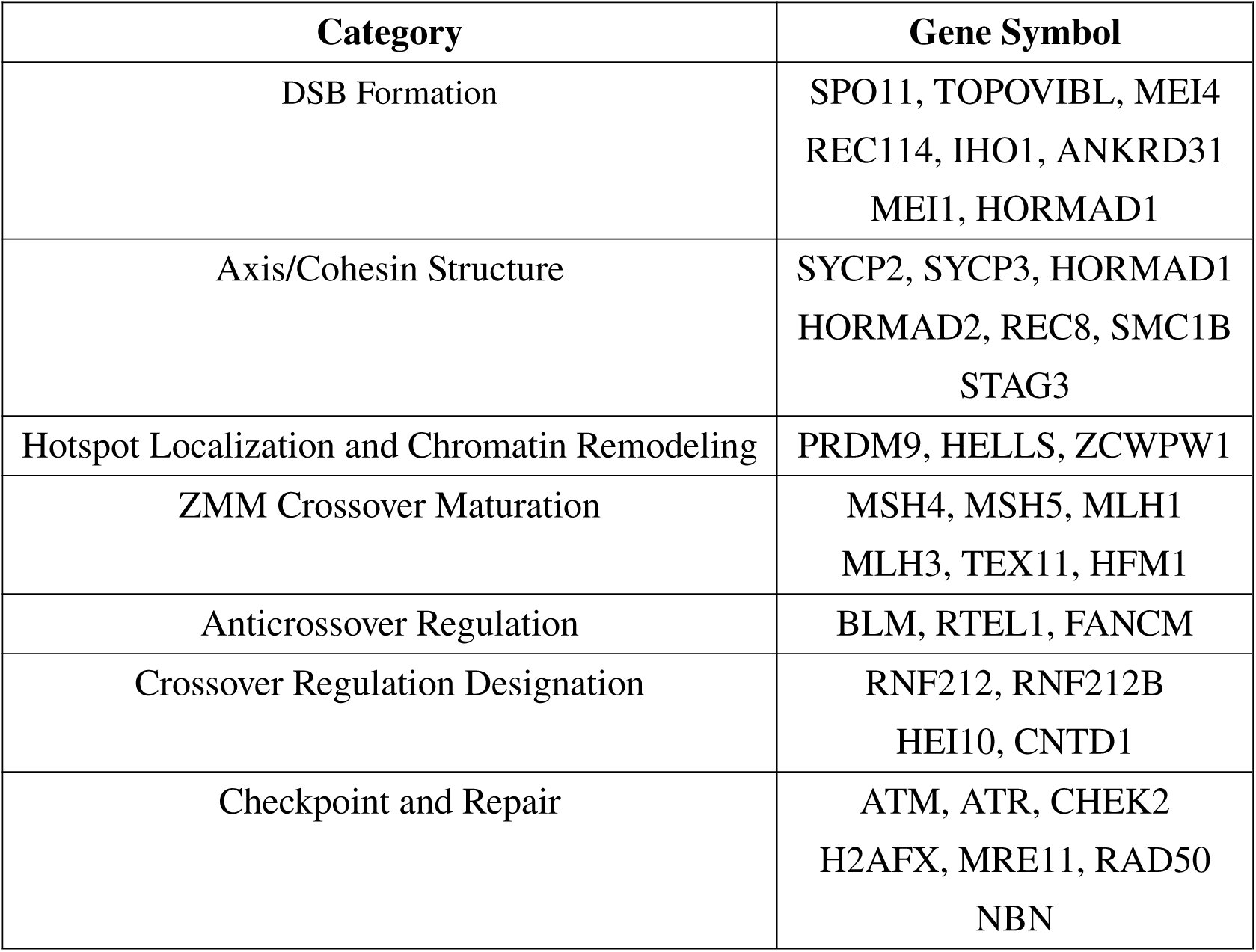
Gene enrichment sets. We apply STRING gene interaction networks to a list of genes that are known to encode proteins involved in meiotic recombination or closely related processes in order to find their interacting neighbors that were 2 hops away, and annotated SNPs to the genomic locations for these genes, split into categories. The categories and gene symbols are given in the table.

**Table S12:** Significant SNPs found in a fixed effects model correcting CO for age. Chromosome number (chr), rsid identifier (SNP), basepair position and the significance of each SNP that is associated in the meta-analysis in fixed effects model that fits age, SNP, and a term to correct for repeated measures, for females and males in MoBa, EstBB and GS. ’na’ indicates that the SNP is not available in this dataset. ’-’ refers to a p-value above 0.05, ’*’ p-value between 0.01 and 0.05, ’**’ to p-value between 0.01 and 0.001 and ’***’ to p-value below 0.001.

| chr | SNP | position | MoBa |  | EstBB |  |
| --- | --- | --- | --- | --- | --- | --- |
|  |  |  | female | male | female | male |
| 1 | rs277680 | 6784464 | - | - | * | - |
| 1 | rs6686883 | 76259146 | na | na | - | - |
| 2 | rs10205866 | 54493250 | * | - | *** | - |
| 3 | rs1799977 | 37053568 | * | - | *** | - |
| 4 | rs3816474 | 1050653 | *** | *** | *** | *** |
| 4 | rs2290409 | 1067249 | *** | *** | - | *** |
| 4 | rs59831071 | 113607978 | - | * | - | ** |
| 6 | rs9467736 | 26380032 | *** | - | *** | - |
| 10 | rs8192775 | 135348026 | - | ** | * | ** |
| 14 | rs1953549 | 20760066 | *** | - | *** | - |
| 14 | rs1254251 | 60818850 | *** | ** | *** | *** |
| 14 | rs1961540 | 91894792 | *** | - | *** | - |
| 16 | rs1623559 | 1858805 | - | ** | ** | - |
| 16 | rs7186699 | 75005197 | - | - | - | - |
| 17 | rs3785884 | 44057595 | *** | - | *** | - |
| 18 | rs1474043 | 12951738 | * | - | *** | - |
| 18 | rs581504 | 74643176 | ** | - | * | - |
| 19 | rs34275421 | 10644812 | * | - | *** | * |
| 19 | rs1230295 | 22621678 | ** | - | * | - |
| 20 | rs450739 | 1210647 | - | - | - | ** |
| 20 | rs4081327 | 61327966 | - | - | ** | - |
| 22 | rs9626333 | 45831225 | *** | - | *** | - |

**Table S13:**
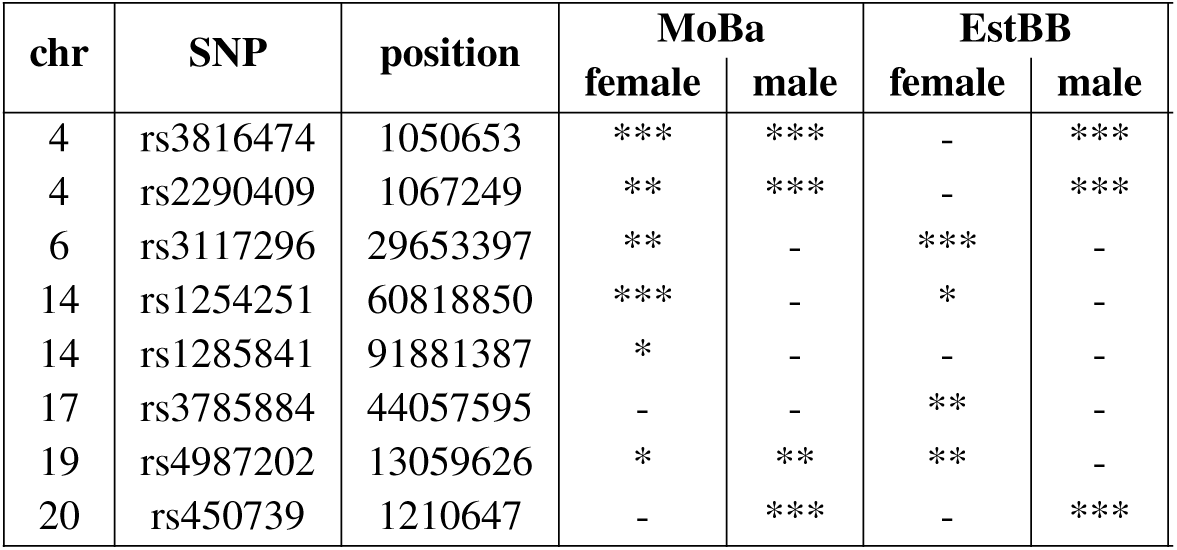
Significant SNPs found in a fixed effects model correcting PA for age. Chromosome number (chr), rsid identifier (SNP), basepair position, and the significance of each SNP that is associated in the meta-analysis in fixed effects model that fits age, SNP, and a term to correct for repeated measures, for females and males in MoBa and EstBB. ’na’ indicates that the SNP is not available in this dataset. ’-’ refers to a p-value above 0.05, ’*’ p-value between 0.01 and 0.05, ’**’ to p-value between 0.01 and 0.001 and ’***’ to p-value below 0.001.

**Table S14:** Significant SNPs found in a fixed effects model correcting TD for age. Chromosome number (chr), rsid identifier (SNP), basepair position, and the significance of each SNP that is associated in the meta-analysis in fixed effects model that fits age, SNP, and a term to correct for repeated measures, for females and males in MoBa and EstBB. ’na’ indicates that the SNP is not available in this dataset. ’-’ refers to a p-value above 0.05, ’*’ p-value between 0.01 and 0.05, ’**’ to p-value between 0.01 and 0.001 and ’***’ to p-value below 0.001.

| chr | SNP | position | MoBa |  | EstBB |  |
| --- | --- | --- | --- | --- | --- | --- |
|  |  |  | female | male | female | male |
| 2 | rs10205866 | 54493250 | * | - | *** | * |
| 4 | rs2290409 | 1067249 | - | *** | - | * |
| 5 | rs143073807 | 122535307 | na | na | ** | - |
| 12 | rs36179555 | 133802299 | ** | - | ** | - |
| 14 | rs2781366 | 20828484 | ** | ** | - | - |
| 14 | rs8015263 | 60958880 | - | - | - | - |
| 14 | rs1961540 | 91894792 | ** | - | *** | - |
| 17 | rs3785884 | 44057595 | *** | * | *** | - |
| 19 | rs4987202 | 13059626 | *** | * | * | - |
| 20 | rs450739 | 1210647 | - | *** | - | * |

**Table S15:**
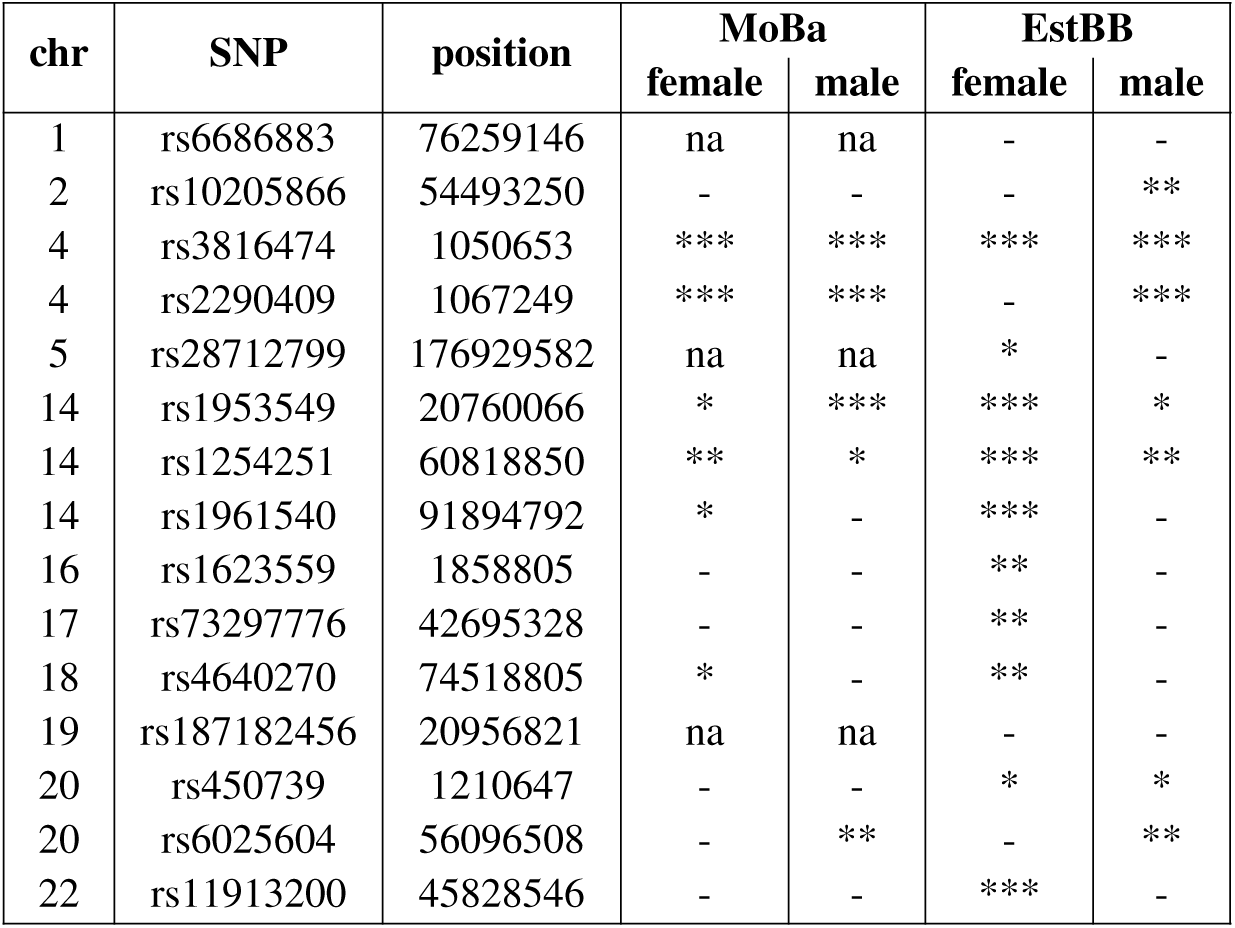
Significant SNPs found in a fixed effects model correcting MD for age. Chromosome number (chr), rsid identifier (SNP), basepair position, and the significance of each SNP that is associated in the meta-analysis in fixed effects model that fits age, SNP, and a term to correct for repeated measures, for females and males in MoBa and EstBB. ’na’ indicates that the SNP is not available in this dataset. ’-’ refers to a p-value above 0.05, ’*’ p-value between 0.01 and 0.05, ’**’ to p-value between 0.01 and 0.001 and ’***’ to p-value below 0.001.

## References and Notes

1. K. Brick, et al., Extensive sex differences at the initiation of genetic recombination. Nature 561 (7723), 338–342 (2018).

2. G. Palsson, et al., Complete human recombination maps. Nature 639 (8055), 700–707 (2025).

3. S. M. Fielder, R. Kempfer, W. G. Kelly, Multiple Sex-Specific Differences in the Regulation of Meiotic Progression in C. elegans. bioRxiv (2020), doi:10.1101/2020.03.12.989418.

4. J. M. Sardell, M. Kirkpatrick, Sex differences in the recombination landscape. Am. Nat. 195 (2), 361–379 (2020).

5. C. Bhérer, C. L. Campbell , A. Auton, Refined genetic maps reveal sexual dimorphism in human meiotic recombination at multiple scales. Nature Communications 8 (1), 14994 (2017).

6. L. Ollivier, B. Charlesworth, F. Pouyet, Beyond recombination: Exploring the impact of meiotic frequency on genome-wide genetic diversity. PLoS Genet. 21 (8), e1011798 (2025).

7. S. E. Johnston, Understanding the genetic basis of variation in meiotic recombination: Past, present, and future. Mol. Biol. Evol. 41 (7) (2024).

8. A. Kong, et al., Common and low-frequency variants associated with genome-wide recombi-nation rate. Nature Genetics 46 (1), 11–16 (2014).

9. C. L. Campbell, N. A. Furlotte, N. Eriksson, D. Hinds, A. Auton, Escape from crossover interference increases with maternal age. Nat. Commun. 6 (1), 6260 (2015).

10. B. V. Halldorsson, et al., Characterizing mutagenic effects of recombination through a sequence-level genetic map. Science 363 (6425), eaau1043 (2019).

11. R. E. Brandlistuen, et al., Cohort Profile Update: The Norwegian Mother, Father and Child Cohort (MoBa). International Journal of Epidemiology 54 (5) (2025).

12. R. J. Hofmeister, et al., Parent-of-origin effects on complex traits in up to 236,781 individuals. Nature 646 (8085), 647–656 (2025).

13. C. Veller, N. Kleckner, M. A. Nowak, A rigorous measure of genome-wide genetic shuffling that takes into account crossover positions and Mendel’s second law. Proc. Natl. Acad. Sci. U. S. A. 116 (5), 1659–1668 (2019).

14. K. Yuan, et al., Fine-mapping across diverse ancestries drives the discovery of putative causal variants underlying human complex traits and diseases. Nat. Genet. 56 (9), 1841–1850 (2024).

15. K. Watanabe, E. Taskesen, A. van Bochoven, D. Posthuma, Functional mapping and annotation of genetic associations with FUMA. Nat. Commun. 8 (1) (2017).

16. A. D. Bell, et al., Insights into variation in meiosis from 31,228 human sperm genomes. Nature 583 (7815), 259–264 (2020).

17. S. A. Carioscia, et al., Common variation in meiosis genes shapes human recombination and aneuploidy. Nature (2026), doi:10.1038/s41586-025-09964-2, 10.1038/s41586-025-09964-2.

18. J. Dai, et al., Molecular basis of the dual role of the Mlh1-Mlh3 endonuclease in MMR and in meiotic crossover formation. Proc. Natl. Acad. Sci. U. S. A. 118 (23), e2022704118 (2021).

19. X. Lin, et al., An in-frame deletion mutation in MLH1 drives Lynch syndrome-associated colorectal cancer. BMC Cancer 25 (1), 1165 (2025).

20. F. Di Tullio, M. Schwarz, H. Zorgati, S. Mzoughi, E. Guccione, The duality of PRDM proteins: epigenetic and structural perspectives. FEBS J. 289 (5), 1256–1275 (2022).

21. K. Farrell, K. Uh, K. Lee, 45 Expression patterns of PRDM family genes in porcine pre-implantation embryos. Reprod. Fertil. Dev. 32 (2), 148–148 (2019).

22. Z. Xu, et al., Pan-cancer analysis identified DOK3 as a novel biomarker for predicting prognosis and immunotherapy effectiveness. *Discov*. Oncol. 16 (1), 2290 (2025).

23. T. L. Peoples-Holst, S. M. Burgess, Multiple branches of the meiotic recombination pathway contribute independently to homolog pairing and stable juxtaposition during meiosis in budding yeast. Genes Dev. 19 (7), 863–874 (2005).

24. E. Anton, C. Zurera-Egea, R. Farriol, Z. Sarrate, J. Blanco, Systematic review and evidence-based classification of differentially expressed mRNA in human spermatozoa: insights to improve the diagnosis and prognosis of male infertility. Reprod. Biomed. Online 51 (3), 104993 (2025).

25. S. Kim, J. A. Govindan, Z. J. Tu, D. Greenstein, SACY-1 DEAD-Box helicase links the somatic control of oocyte meiotic maturation to the sperm-to-oocyte switch and gamete maintenance in Caenorhabditis elegans. Genetics 192 (3), 905–928 (2012).

26. S. Shinriki, et al., DDX41 coordinates RNA splicing and transcriptional elongation to prevent DNA replication stress in hematopoietic cells. Leukemia 36 (11), 2605–2620 (2022).

27. Y. Wei, et al., TORC1 regulators Iml1/GATOR1 and GATOR2 control meiotic entry and oocyte development in Drosophila. Proc. Natl. Acad. Sci. U. S. A. 111 (52), E5670–7 (2014).

28. C. M. Lake, R. J. Nielsen, R. S. Hawley, The Drosophila zinc finger protein trade embargo is required for double strand break formation in meiosis. PLoS Genet. 7 (2), e1002005 (2011).

29. J. K. Singh, H. van Attikum, DNA double-strand break repair: Putting zinc fingers on the sore spot. Semin. Cell Dev. Biol. 113, 65–74 (2021).

30. L. Mukhametzyanova, et al., Activation of recombinases at specific DNA loci by zinc-finger domain insertions. Nat. Biotechnol. 42 (12), 1844–1854 (2024).

31. C. M. Phillips, A. F. Dernburg, A family of zinc-finger proteins is required for chromosome-specific pairing and synapsis during meiosis in C. elegans. Dev. Cell 11 (6), 817–829 (2006).

32. A. W. Y. Leung, et al., 3’-Phosphoadenosine 5’-phosphosulfate synthase 1 (PAPSS1) knock-down sensitizes non-small cell lung cancer cells to DNA damaging agents. Oncotarget 6 (19), 17161–17177 (2015).

33. I. Krätschmer, et al., Discovery of shared epigenetic pathways across human phenotypes. bioRxiv (2024), doi:doi:10.1101/2024.04.15.589547.

34. Z. Zheng, et al., Leveraging functional genomic annotations and genome coverage to improve polygenic prediction of complex traits within and between ancestries. Nat. Genet. 56 (5), 767–777 (2024).

35. D. Speed, D. J. Balding, SumHer better estimates the SNP heritability of complex traits from summary statistics. Nat. Genet. 51 (2), 277–284 (2019).

36. A. Hochwagen, G. A. Marais, Meiosis: A PRDM9 Guide to the Hotspots of Recombination. Current Biology 20 (6), R271–R274 (2010).

37. C. Girard, D. Zwicker, R. Mercier, The regulation of meiotic crossover distribution: a coarse solution to a century-old mystery? Biochem. Soc. Trans. 51 (3), 1179–1190 (2023).

38. B. Alleva, K. Brick, F. Pratto, M. Huang, R. D. Camerini-Otero, Cataloging human PRDM9 Allelic variation using long-read sequencing reveals PRDM9 population specificity and two distinct groupings of related alleles. Front. Cell Dev. Biol. 9, 675286 (2021).

39. A. Kong, et al., Recombination rate and reproductive success in humans. Nat. Genet. 36 (11), 1203–1206 (2004).

40. H. C. Martin, et al., Multicohort analysis of the maternal age effect on recombination. Nat. Commun. 6 (1), 7846 (2015).

41. S. Stankovic, et al., Genetic links between ovarian ageing, cancer risk and de novo mutation rates. Nature 633 (8030), 608–614 (2024).

42. K. S. Rønningen, et al., The biobank of the Norwegian mother and child cohort Study: A resource for the next 100 years. European Journal of Epidemiology 21 (8), 619–625 (2006).

43. E. C. Corfield, et al., The Norwegian Mother, Father, and Child cohort study (MoBa) genotyping data resource: MoBaPsychGen pipeline v.1. bioRxiv (2024), doi:10.1101/2022.06.23.496289.

44. R. J. Hofmeister, et al., Parent-of-Origin inference for biobanks. Nature Communications 13 (6668) (2022).

45. R. J. Hofmeister, D. M. Ribeiro, S. Rubinacci, O. Delaneau, Accurate rare variant phasing of whole-genome and whole-exome sequencing data in the UK Biobank. Nature Genetics 55 (7), 1243–1249 (2023).

46. S. McCarthy, et al., A reference panel of 64,976 haplotypes for genotype imputation. Nature Genetics 48 (10), 1279–1283 (2016).

47. J. Yang, et al., Conditional and joint multiple-SNP analysis of GWAS summary statistics identifies additional variants influencing complex traits. Nat. Genet. 44 (4), 369–75, S1–3 (2012).

48. J. C.-C. Chan, I. Jeliazkov, MCMC Estimation of Restricted Covariance Matrices. Journal of Computational and Graphical Statistics 18 (2), 457–480 (2009), doi:{}.

49. D. Speed, D. J. Balding, SumHer better estimates the SNP heritability of complex traits from summary statistics. Nature Genetics 51 (2), 277–284 (2018).

50. E. J. Orliac, et al., Improving GWAS discovery and genomic prediction accuracy in biobank data. Proc. Natl. Acad. Sci. U. S. A. 119 (31), e2121279119 (2022).

51. D. Szklarczyk, et al., The STRING database in 2023: protein-protein association networks and functional enrichment analyses for any sequenced genome of interest. Nucleic Acids Res. 51 (D1), D638–D646 (2023).

52. B. J. Lesch, S. J. Silber, J. R. McCarrey, D. C. Page, Parallel evolution of male germline epigenetic poising and somatic development in animals. Nat. Genet. 48 (8), 888–894 (2016).

53. J. Guo, et al., The adult human testis transcriptional cell atlas. Cell Res. 28 (12), 1141–1157 (2018).

54. W. Xia, et al., Resetting histone modifications during human parental-to-zygotic transition. Science 365 (6451), 353–360 (2019).

55. N. Altemose, et al., A map of human PRDM9 binding provides evidence for novel behaviors of PRDM9 and other zinc-finger proteins in meiosis. Elife 6 (e28383) (2017).

56. 1000 Genomes Project Consortium, et al., A global reference for human genetic variation. Nature 526 (7571), 68–74 (2015).

57. T. Schoeler, et al., Participation bias in the UK Biobank distorts genetic associations and downstream analyses. *Nat*. Hum. Behav. 7 (7), 1216–1227 (2023).

58. J. O’Connell, et al., A general approach for haplotype phasing across the full spectrum of relatedness. PLoS Genet. 10 (4), e1004234 (2014).

59. G. R. Nagy, B. Györffy, B. Nagy, J. Rigó, Lower risk for Down syndrome associated with longer oral contraceptive use: a case–control study of women of advanced maternal age presenting for prenatal diagnosis. Contraception 87 (4), 455–458 (2013).

60. N. E. Lamb, K. Yu, J. Shaffer, E. Feingold, S. L. Sherman, Association between maternal age and meiotic recombination for trisomy 21. Am. J. Hum. Genet. 76 (1), 91–99 (2005).

61. S. Linn, B. Murtaugh, E. Casey, Role of sex hormones in the development of osteoarthritis. PM R 4 (5 Suppl), S169–73 (2012).

62. X. Tan, et al., Causal association of menstrual reproductive factors on the risk of osteoarthritis: A univariate and multivariate Mendelian randomization study. PLoS One 19 (8), e0307958 (2024).

63. Y. V. Louwers, J. A. Visser, Shared genetics between age at menopause, early menopause, POI and other traits. Front. Genet. 12, 676546 (2021).

